# Cognitive capacity limits are remediated by practice-induced plasticity between the Putamen and Pre-Supplementary Motor Area

**DOI:** 10.1101/564450

**Authors:** K.G. Garner, M.I. Garrido, P.E. Dux

**Affiliations:** Queensland Brain Institute, University of Queensland, Brisbane, Australia; School of Psychology, University of Queensland, Brisbane, Australia; Australian Research Council Centre of Excellence for Integrative Brain Function, Australia; Department of Psychology, University of Birmingham, UK; Melbourne School of Psychological Sciences, University of Melbourne, Melbourne, Australia; Centre for Advanced Imaging, University of Queensland, Brisbane, Australia

**Author notes:** denotes senior authorship (M.I. Garrido, P.E. Dux). **Correspondence to:** K.G. Garner, Queensland Brain Institute, 79, University of Queensland, St. Lucia, QLD, 4072. The authors declare no competing interests. Analysis code: https://github.com/kel-github/multi-practice-repository. Raw data: https://espace.library.uq.edu.au/view/UQ:370251. Processed data: To access the processed data please.

## Abstract

Humans show striking limitations in information processing when multitasking, yet can modify these limits with practice. Such limitations have been linked to a frontal-parietal network, but recent models of decision-making implicate a striatal-cortical network. We adjudicated these accounts by investigating the circuitry underpinning multitasking in 100 individuals and the plasticity caused by practice. We observed that multitasking costs, and their practice induced remediation, are best explained by modulations in information transfer between the striatum and the cortical areas that represent stimulus-response mappings. Specifically, our results support the view that multitasking stems at least in part from taxation in information sharing between the putamen and pre-supplementary motor area (pre-SMA). Moreover, we propose that modulations to information transfer between these two regions leads to practice-induced improvements in multitasking.

**Significance statement:** Humans show striking limitations in information processing when multitasking, yet can modify these limits with practice. Such limitations have been linked to a frontal-parietal network, but recent models of decision-making implicate a striatal-cortical network. We adjudicated these accounts by investigating the circuitry underpinning multitasking in 100 individuals and the plasticity caused by practice. Our results support the view that multitasking stems at least in part from taxation in information sharing between the putamen and pre-supplementary motor area (pre-SMA). We therefore show that models of cognitive capacity limits must consider how subcortical and cortical structures interface to produce cognitive behaviours, and we propose a novel neurophysiological substrate of multitasking limitations.

---

Although human information processing is fundamentally limited, the points at which task difficulty or complexity incurs performance costs are malleable with practice. For example, practicing component tasks reduces the response slowing that is typically induced as a consequence of attempting to complete the same tasks concurrently (multitasking) (Telford, 1931; Ruthruff, Johnston and Van Selst, 2001; Strobach and Torsten, 2017). These limitations are currently attributed to competition for representation in a frontal-parietal network (Watanabe and Funahashi, 2014, 2018; Garner and Dux, 2015; Marti, King and Dehaene, 2015), in which the constituent neurons adapt response properties in order to represent the contents of the current cognitive episode (Duncan, 2010, 2013; Woolgar *et al.*, 2011). Despite recent advances, our understanding of the network dynamics that drive multitasking costs and the influence of practice remains unknown. Furthermore, although recent work has focused on understanding cortical contributions to multitasking limitations, multiple theoretical models implicate striatal-cortical circuits as important neurophysiological substrates for the execution of single sensorimotor decisions (Joel, Niv and Ruppin, 2002; Bornstein and Daw, 2011; Caballero, Humphries and Gurney, 2018), the formation of stimulus-response representations in frontal-parietal cortex (Ashby, Turner and Horvitz, 2010; Hélie, Ell and Ashby, 2015), and performance of both effortful and habitual sensorimotor tasks (Yin and Knowlton, 2006; Graybiel and Grafton, 2015; Jahanshahi *et al.*, 2015). This suggests that a complete account of cognitive limitations and their practice-induced attenuation also requires interrogation into the contribution of striatal-cortical circuits. We seek to address these gaps in understanding by investigating how multitasking and practice influence network dynamics between striatal and cortical regions previously implicated in the cognitive limitations that give rise to multitasking costs (Garner and Dux, 2015).

We previously observed that improvements in the decodability of component tasks in two regions of the frontal-parietal network - pre-supplementary motor area (pre-SMA/SMA), the intraparietal sulcus (IPS) - and one region of the striatum (putamen) predicted practice-induced multitasking improvements (Garner and Dux, 2015). This implies that practice may not divert performance from the frontal-parietal system, as had been previously assumed (Petersen *et al.*, 1998; Kelly and Garavan, 2005; Yin and Knowlton, 2006; Chein and Schneider, 2012), but rather, may alleviate multitasking costs by reducing competition for representation within the same system. Moreover, our finding that the putamen showed changes to task decodability that predicted behavioural improvements comparable to what was observed for pre-SMA and IPS implies that rather than stemming from overload of an entirely cortical network (Marois and Ivanoff, 2005; Dux *et al.*, 2006; Marti, King and Dehaene, 2015), multitasking costs are manifest by limitations within a distributed striatal-cortical system. This raises the question of how interactions between these brain regions give rise to multitasking costs and how can these be mitigated with practice: Do multitasking costs reflect over-taxation of striatal-cortical circuits? Or are they a consequence of competition for representation between cortical areas? Alternately, do multitasking costs stem from limitations in both striatal-cortical and corticocortical connections? Does practice alleviate multitasking costs via modulating all the interregional couplings that give rise to multitasking behaviour, or by selectively increasing or reducing couplings between specific regions?

Our aim was to arbitrate between these possibilities by applying dynamic causal modelling (DCM, Friston, Harrison and Penny, 2003) to an fMRI dataset (N=100) collected while participants performed a multitasking paradigm before and after practice on the same paradigm (N=50) or on an active control task (N=50) (Garner and Dux, 2015). We sought to first characterise the modulatory influence of multitasking on the network dynamics between the pre-SMA, IPS and putamen, and then to understand how practice modulated these network dynamics to drive multitasking performance improvements.

## Methods

### Participants

The MRI scans of the participants (N=100) previously analysed in (Garner and Dux, 2015) were included in the present analysis, apart from the data of 2 participants, for whom some of the scans were corrupted due to experimenter error. The remaining 98 participants had been pseudorandomly allocated to the practice group (N=48, mean age: 24.33 [sd: 6.31], 44 F, 44 right handed) or the control group (N=50, mean age: 24.58 [sd: 5.48], 46 F, 45 right-handed). All participants received 10 AUD per hour for participation. Participants also earned bonus dollars across the three training sessions. Bonus dollars were accrued for high accuracy and for beating RT deadlines (~20 AUD per participant). For details of the original data point exclusions, we refer the reader to Garner and Dux (2015). The University of Queensland Human Research Ethics Committee approved the study as being within the guidelines of the National Statement on Ethical Conduct in Human Research and all participants gave informed, written consent.

### Experimental Protocols

Participants attended six experimental sessions: a familiarization session, two MRI sessions and three behavioural practice sessions. Familiarization sessions were conducted the Friday prior to the week of participation, where participants learned the stimulus-response mappings and completed two short runs of the task. The MRI sessions were conducted to obtain pre-practice (Monday session) and post-practice (Friday session) measures. These sessions were held at the same time of day for each participant. Between the two MRI sessions, participants completed three behavioural practice sessions, where they either practiced the multitasking paradigm (practice group) or the visual-search task (control group). Participants typically completed one practice session per day, although on occasion two practice sessions were held on the same day to accommodate participants’ schedules (when this occurred, the two sessions were administered with a minimum of an hour break between them). Participants also completed an online battery of questionnaires that formed part of a different study.

#### Behavioural Tasks

All tasks were programmed using Matlab R2010a (Mathworks, Natick, MA) and the Psychophysics Toolbox v3.0.9 extension (23). The familiarization and behavioural training sessions were conducted with a 21-inch, Sony Trinitron CRT monitor and a Macintosh 2.5 GHz Mini computer.

#### Multitasking Paradigm

For each trial of the multitasking paradigm, participants performed either one (single-task condition) or two (multitask condition) sensorimotor tasks. Both involved a 2-alternative discrimination (2-AD), mapping the two stimuli to two responses. For one task, participants were presented with one of two white shapes that were distinguishable in terms of their smooth or spikey texture, presented on a black screen and subtending approximately 6° of visual angle. The shapes were created using digital sculpting software (Scluptris Alpha 6) and Photoshop CS6. Participants were required to make the appropriate manual button press to the presented shape, using either the index or middle finger of either the left or right hand (task/hand assignment was counterbalanced across participants). For the other task, participants responded to one of two complex tones using the index or middle finger of the hand that was not assigned to the shape task. The sounds were selected to be easily discriminable from one another. Across both the single-task and multitask trial types, stimuli were presented for 200 ms, and on multitask trials, were presented simultaneously.

#### Familiarisation Session

During the familiarization session, participants completed two runs of the experimental task. Task runs consisted of 18 trials, divided equally between the three trial types (shape single-task, sound single-task, and multitask trials). The order of trial type presentation was pseudo-randomised. The first run had a *short inter-trial-interval (ITI)* and the trial structure was as follows; an alerting fixation dot, subtending 0.5° of visual angle was presented for 400 ms, followed by the stimulus/stimuli that was presented for 200 ms. Subsequently a smaller fixation dot, subtending 0.25° of visual angle, was presented for 1800 ms, during which participants were required to respond. Participants were instructed to respond as accurately and quickly as possible to all tasks. For the familiarization session only, performance feedback was then presented until the participant hit the spacebar in order to continue the task. For example, if the participant had completed the shape task correctly, they were presented with the message ‘You got the shape task right’. If they performed the task incorrectly, the message ‘Oh no! You got the shape task wrong’ was displayed. On multitask trials; feedback was presented for both tasks. If participants failed to achieve at least 5/6 trials correct for each trial type they repeated the run until this level of accuracy was attained.

The second run familiarized participants with the timing of the paradigm to be used during the MRI sessions - a slow event-related design with a *long ITI*. The alerting fixation was presented for 2000 ms, followed by the 200 ms stimulus presentation, 1800 ms response period and feedback. Subsequently an ITI, during which the smaller fixation dot remained on screen, was presented for 12000 ms.

#### MRI Sessions

Participants completed six long ITI runs in the scanner, with 18 trials per run (6 of each trial type, pseudo-randomly ordered for each run), for a total of 108 trials for the session. Trial presentation was identical to the long ITI run presented at the familiarization session, except that feedback was not presented at the end of each trial.

#### Practice Sessions

All participants were informed that they were participating in a study examining how practice improves attention, with the intention that both the practice and control groups would expect their practice regimen to improve performance. The first practice session began with an overview of the goals of the practice regimen; participants were informed that they were required to decrease their response time (RT), while maintaining a high level of accuracy. The second and third sessions began with visual feedback in the form of a line graph, plotting RT performance from the previous practice sessions.

For each session, participants completed 56 blocks of 18 trials, for a total of 1008 trials, resulting in 3024 practice trials overall. To ensure that participants retained familiarity with the timings of the task as presented in the scanner, between 2 and 4 of the blocks in each session used long ITI timings.

The practice group performed the multitasking paradigm, as described above (see Familiarization Session), except that performance feedback was not displayed after each trial. Over the course of practice, participants from this group performed 1008 trials of each trial type (shape single-task, sound single-task, multitask). Participants in the control group went through the identical procedures to the practice group, except that they completed a visual search task instead of the multitasking paradigm. Participants searched for a ‘T’ target amongst 7, 11, or 15 rotated ‘L’s’ (to either 90° or 270°). Participants indicated whether the target was oriented to 90° or 270°, using the first two fingers of their left or right hand (depending upon handedness). Over the course of the three practice sessions, participants completed 1008 trials for each set size.

For both groups performance feedback showed mean RT (collapsed across the two single-tasks for the practice group, and over the three set-sizes for the control group), and accuracy, for the previous 8 blocks, total points scored, and the RT target for the subsequent 8 blocks. If participants met their RT target for over 90 % of trials, and achieved greater than 90 % accuracy, a new RT target was calculated by taking the 75^th^ percentile of response times recorded over the previous 8 blocks. Furthermore, 2 points were awarded. If participants did not beat their RT target for over 90 % trials, but did maintain greater than 90 % accuracy, 1 point was awarded.

#### MRI Data Acquisition

Images were acquired using a 3T Siemens Trio MRI scanner (Erlangen, Germany) housed at the Centre for Advanced Imaging at The University of Queensland. Participants lay supine in the scanner and viewed the visual display via rear projection onto a mirror mounted on a 12-channel head coil. A T1-weighted anatomic image was collected after the fourth experimental run of the scanning session (repetition time (TR) = 1.9 s, echo time (TE) = 2.32 ms, flip angle (FA) = 9°, field of view (FOV) = 192 × 230 × 256 mm, resolution = 1 mm^3^). Functional T2*-weighted images were acquired parallel to the anterior commissure-posterior commissure plane using a GRE EPI sequence (TR = 2 s, TE = 35 ms, FA = 79 °, FOV = 192 × 192 mm, matrix = 64 × 64, in-plane resolution = 3 × 3 mm). Each volume consisted of 29 slices (thickness = 3 mm, interslice gap = .5 mm), providing whole brain coverage. We synchronized the stimulus presentation with the acquisition of functional volumes.

#### MRI Data Analysis

fMRI data were preprocessed using the SPM12 software package (Wellcome Trust Centre for Neuroimaging, London, UK; http://www.fil.ion.ucl.ac.uk/spm). Scans from each subject were corrected for slice timing differences using the middle scan as a reference, realigned using the middle first as a reference, co-registered to the T1 image, spatially normalised into MNI standard space, and smoothed with a Gaussian kernel of 8 mm full-width at half maximum.

#### Dynamic Causal modelling

To assess the causal direction of information flow between brain regions, we applied Dynamic Causal Modelling (DCM), which maps experimental inputs to the observed fMRI output, via hypothesised modulations to neuronal states that are characterised using a biophysically informed generative model (Friston, Harrison and Penny, 2003). Parameter estimates are expressed as rate constants (i.e. the rate of change of gross neural activity in one region, given the activity in the coupled brain region), and are fit using Bayesian parameter estimation. It is important to note that any interpretations regarding information transfer between brain regions is based on the assumption that these rate parameters reflect a causal relationship between the regions of interest that is meaningful with regards to task performance. Moreover, with DCM, we seek to model coupling changes between regions of interest that have been defined a priori, and that the presence of such inter-regional couplings are postulated. It is possible that any observed coupling changes could be driven by a third node that is not included in the proposed network architecture. Moreover, the currently proposed architectures certainly do not reflect the entire network that underpins multitasking of sensorimotor tasks.

#### DCM Implementation

Implementation of DCM requires definition of endogenous connections within the network (A parameters), the modulatory influence of experimental factors (B parameters), and the influence of exogenous/driving inputs into the system (e.g. sensory inputs, C parameters) (Friston, Harrison and Penny, 2003). We implemented separate DCM’s to investigate i) the modulatory influence of multitasking on the pre-practice data, and ii) the modulatory influence of practice on the pre- to post-practice data.

To make inferences regarding the modulatory influence of multitasking, we defined our endogenous network as comprising reciprocal connectivity between all three of our ROIs, on the basis of anatomical and functional evidence for connections between all three of them (Alexander, DeLong and Strick, 1986; Cavada and Goldman-Rakic, 1989; Luppino *et al.*, 1993; Wise *et al.*, 1997; Haber, 2016). To address our theoretically motivated question regarding the locus of modulatory influence of the multitasking, we first implemented all 63 possible combinations of the modulatory influence of the multitasking (i.e. allowing each combination of connections to be modulated by the multitasking factor, see Extended Figure 2B for an illustration of the model architectures) and then grouped the model space into 3 families: those that allowed any combination of corticocortical modulations, but not striatal-cortical (*corticocortical family,* with 3 models in total M_1-3_ = 3), those that allowed the reverse pattern (*striatal-cortical family,* with 15 models in total, M_4-18_, and those that allowed modulations to both types of connections (*both family*, with 45 models in total, M_19-63_). We found it very difficult to define the most likely locus of input *a priori*, given empirical evidence that both the striatum and the intraparietal sulcus receive inputs from sensory pathways (Saint-Cyr, Ungerleider and Desimone, 1990; Grefkes and Fink, 2005; Anderson *et al.*, 2010; Reig and Silberberg, 2014; Vossel, Geng and Fink, 2014; Alloway *et al.*, 2017; Guo *et al.*, 2018). Instead we opted to first determine whether the data were better modelled using the putamen or the IPS as the input. Importantly, this parameter did not vary over experimental conditions, therefore, this parameter did not explain changes in network activity that were attributable to the multitasking manipulation. We therefore implemented the full set of models [M_1-63_] with inputs to either the IPS, or to the putamen, so that we could test which input best explained the data (invariant to whether the input was from a single- or multitask trial). Thus we fit a total of 126 (2×63) models to the pre-practice data.

To make inferences regarding the modulatory influence of practice on both single and multitask conditions, we carried out the following for both the single-task and the multitask data (see below for details on data extraction): based on the endogenous connectivity and locus of driving input identified by the preceding analysis, we then fit the 15 possible modulatory influences of the practice factor (i.e. pre- to post-practice).

#### Extraction of fMRI Signals for DCM Analysis

The brain regions of interest (ROIs) were selected by the following steps: first we identified regions that showed increased activity for both single tasks at the pre-training session, second, we sought which of these showed increased activity for multitask trials relative to single task trials. Lastly, we asked which of these regions also showed a practice (pre vs post) by group interaction (Garner and Dux, 2015). The left and right intraparietal sulcus (IPS), left and right putamen, and the supplementary motor area (SMA) were implicated by this interaction. In the interest of reducing the complexity of the model space, and in the absence of lateralized differences in the current data, we included only regions in the left hemisphere and the SMA in the current analysis.

For each region, we restricted the initial search radius by anatomically defined ROI masks, and extracted the first eigenvariate of all voxels within a sphere of 4 mm radius centered over the participant specific peak for the initial contrast (increased activity for both single tasks, as in the previous study), adjusted for the effects of interest (p < .05, uncorrected). We opted to use this approach, rather than selecting a fixed functional ROI across participants, as we know that there are clear individual differences in the exact peak of BOLD signal changes within the brain regions that constitute the multiple demand network when participants perform comparable sensorimotor tasks (Crittenden and Duncan 2014). Therefore we sought to ensure that we identify the voxels for each participant that are most responsive to the functional localiser of interest, while leveraging *a priori* knowledge gained by the group-level contrasts (N=100) and prior anatomical knowledge. This, in our opinion, utilises a good combination of our a priori knowledge concerning brain structure and function, in order to localise meaningful BOLD signal changes at the individual level.

We created the anatomical masks in standard MNI space using FSL. As can be seen from Extended Figure 2d, participant-level peaks tended to cluster within the anatomically defined region, as would have occurred had we used a spherical ROI based on the functional data. For the IPS we used the Juelich Histological atlas and for the putamen and the SMA we used the Harvard-Oxford cortical and subcortical atlas. Note: to analyse the modulatory influence of practice on single-task data, we regressed out activity attributable to the multitask condition at this step. To analyse the modulatory influence of practice on multitask data, we comparably regressed out the single-task data at this step. For the first analysis concerning the multitasking network, we concatenated the 6 functional pre-training runs to form a single time series, and for the analysis of the influence of practice, we concatenated the 6 pre-training and 6 post-training runs [total runs = 12]. The two resulting time series were each adjusted for confounds using regressors for movement and for each run. The DCMs were fit using the resulting time series and hence provide a global estimate for the interactions amongst areas across the whole experiment rather than trial-specific estimates. It is reasonable to expect that over the course of the experiment there will be some degree of variability between the flow of information from putamen to pre-SMA, due to potential fatigue and/or over learning effects. These nuisance effects are however mitigated in our event-related design.

#### Bayesian Model Comparison and Inference over Parameters

As our hypotheses concerned the modulatory influence of our experimental factors on model characteristics, rather than any specific model *per se*, we implemented random effects bayesian model comparison between model families (Penny *et al.*, 2010), with both family inference and Bayesian model averaging (BMA) as implemented in SPM 12. We opted to apply a random effects approach that uses a hierarchical Bayesian model to estimate the parameters of a Dirichlet distribution over all models, to protect against the distortive influence of outliers (Stephan *et al.*, 2009). Specifically, the Dirichlet density describes the probability of each model, given the probability of all the models across the group. Its parameters can be considered as a proxy for a count for how many times a model won across participants. Therefore, improbable individual contributions to the group-level data are down-weighted proportional to the likelihood of the observation and contribute less to the evidence over the model space (e.g. a model that only wins for one participant will not hold much weight in the model probability space). This therefore mitigates the potential influence of individual outliers on the model selection procedures. For each family comparison we report i) the expectation of the posterior probability (i.e. the expected likelihood of obtaining the model family *k*, given the data p(f_k_|Y)), and ii) the exceedance probability of family *k* being more likely than the alternative family *j*, given the data p(f_k_ > f_j_ |Y), see (Penny *et al.*, 2010)). To ensure that a particular family of models is not advantaged due to merely containing more models than a comparison family, a uniform prior needs to be set at the family level. The prior over a given family is defined as *p*(*f*_*k*_) = 1/*K*, where *K* is the total number of families. As the prior at the family level is obtained by summing the priors across constituent models in the family set, the uniform family prior is implemented at the model ( *m* ) level as *p*(*m*) = 1/*KN*_*k*_ ∀ *m* ∈ *f*_*k*_, where *N* is the number of models in family *k* (Penny et al. 2010).

Upon establishment of the winning family, we sought to identify, post-hoc, which specific parameters were likely, given the data, and when relevant, where there was evidence for group differences. To achieve this, we calculated the posterior probability (Pp) that the posterior density over the given parameter has deviated from zero (or in the case of group differences, whether the difference between posterior estimates has deviated from zero), using the SPM spm_Ncdf.m function. To correct for multiple comparisons, we reported Pp’s as having deviated from zero when the likelihood exceeded that set by the Sidak correction (1 - *α*)^1/m^ where m = the number of null hypotheses being tested.

## Results

As all results unrelated to the dynamic causal modelling analysis are described in detail in Garner and Dux (2015), we recap the relevant findings here. Participants completed a multitasking paradigm (Figure 1a) while being scanned with functional magnetic resonance imaging (fMRI), in a slow event-related design. For the multitasking paradigm, participants completed both single- and multi-task trials. For the single-task trials, participants made a 2-alternative discrimination between either one of two equiprobable shapes (visual-manual task), or between one of two equiprobable sounds (auditory-manual task). Participants were instructed to make the correct button-press as quickly and as accurately as possible. On multitask trials, the shape and sound stimuli were presented simultaneously, and participants were required to make both discriminations (visual-manual task and auditory-manual task) as quickly and as accurately as possible. Between the pre- and post-practice scanning sessions, participants were randomly allocated to a practice group or an active-control group (also referred to as the control group). The practice group performed the multitask paradigm over multiple days whereas the control group practiced a visual-search task (Figure 1b). For both groups, participants were adaptively rewarded for maintaining accuracy while reducing response-time (see methods section for details).

**Figure 1:**
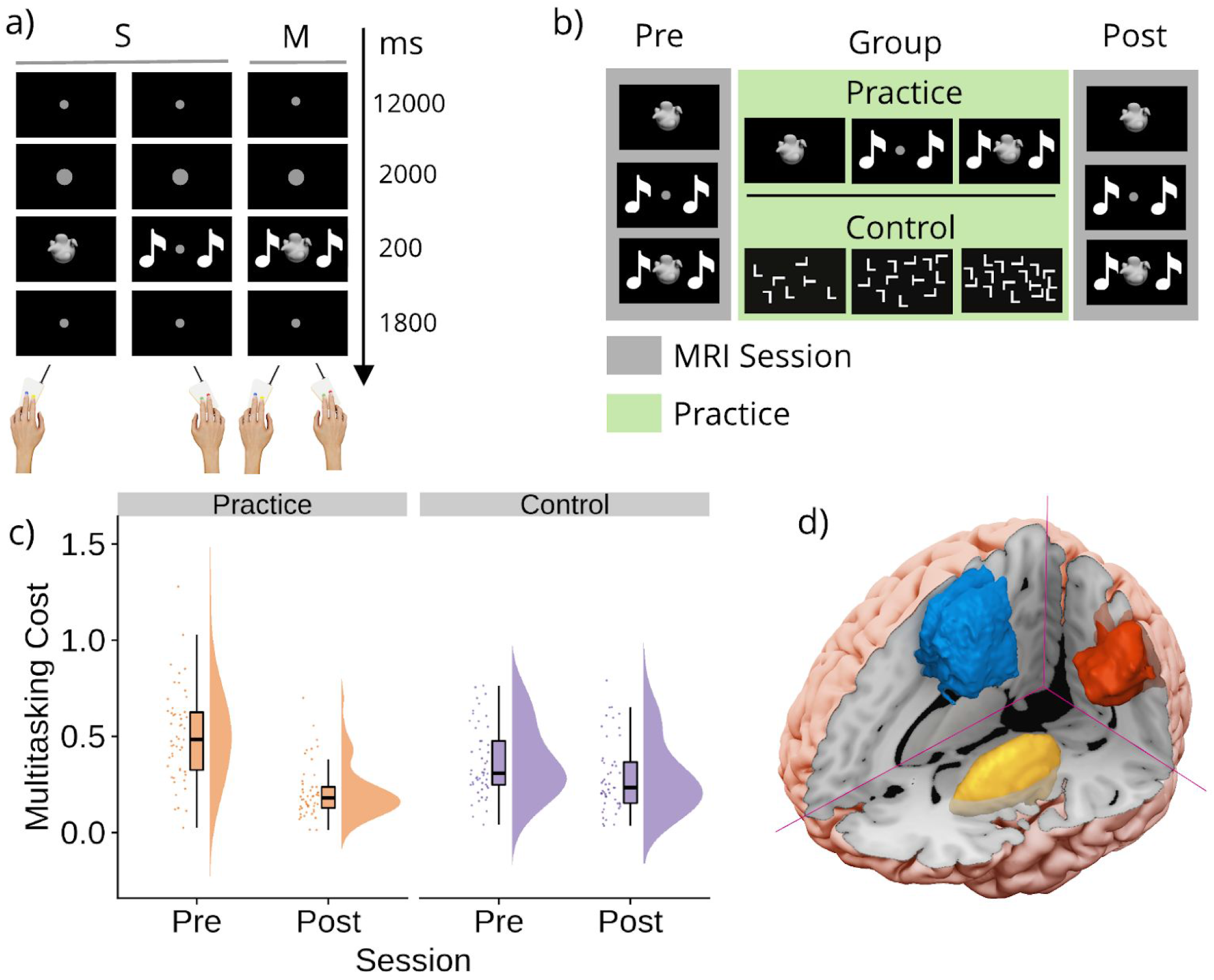
Task, protocol, behaviour and regions of interest. a) Multitasking paradigm: The task was comprised of two single-task (S) conditions and one multitask (M) condition. Each S was a 2 alternative-discrimination between either one of two equiprobable stimuli. The stimuli could either be shapes (visual-manual task), or sounds (auditory-manual task). On M trials, participants were required to complete both Ss (visual-manual and auditory-manual). On all trials, participants were requested to perform the task(s) as quickly and as accurately as possible. b) Protocol: At both the pre- and post- practice sessions, all participants completed the multitasking paradigm while structural and functional MRI images were taken. Participants were then allocated to either the practice or the active-control group. The practice group subsequently performed the multitask paradigm over three sessions, whereas the control group practiced a visual-search task with three levels of difficulty, under a comparable reinforcement regimen. c) Multitasking costs to response time [mean(Ms) - mean(Ss)] for the practice and control groups, at the pre- and post-practice sessions, presented as individual data points, boxplots and densities (raincloud plots, Allen *et al.*, 2018). d) Regions of interest identified by our previous study (Garner and Dux, 2015); the Supplementary Motor Area (blue), the Intraparietal Sulcus (red), and the Putamen (yellow).

Our key behavioural measure of multitasking costs was the difference in response-time (RT) between the single- and multi-task conditions. Performing the component tasks as a multitask increases RT for both tasks, relative to when each is performed alone as a single task. The effectiveness of the paradigm to assess multitasking was confirmed with multitasking costs being clearly observed in the pre-practice session (main effect of condition, single- vs multi-task, F(1, 98) = 688.74, MSE = .026, p<.0001, η_p_^2^ = .88, see extended Figure 1a). Critically, the practice group showed a larger reduction in multitasking costs between the pre- and post-practice sessions than the control group (significant session (pre vs. post) x condition (single-task vs multitask) x group (practice vs control) interaction; F(1, 98) = 31.12, MSE = .01, p < .001, η_p_^2^ = .24, Figure 1c). Specifically, the practice group showed a mean reduction (pre-cost - post-cost) of 293 ms (95% CI [228, 358]) whereas the control group showed a mean reduction of 79 ms (95% CI: [47, 112]). These findings did not appear to be due to a speed/accuracy trade-off as the group x session x condition interaction performed on the accuracy data was not statistically significant (p=.06).

We sought to identify the brain regions that could be part of the multiple demand network that supports performance of both tasks, as our question pertains to how regions that appear to be associated with cognitive control, invariant to the modality of the underlying tasks, interact under conditions of multitasking and multitasking practice. Specifically, regions of interest were defined as those that 1) showed increased activity for both single tasks (i.e. a conjunctive contrast), as could be expected by brain areas containing neurons that adapt to represent the current cognitive episode and brain areas that contribute to, or at least are sensitive to, the performance of both tasks, 2) showed sensitivity to multitasking demands (i.e. increased activity for multitask relative to single-task trials), and 3) showed specificity in response to the training regimen, i.e. showed a group x session interaction (see Garner and Dux, 2015 for details). Thus our regions of interest are sensitive to both single tasks and to the multitasking practice regimen, regardless of laterality and the modality of the underlying single tasks. This criteria isolated the pre-SMA/SMA, the left and right inferior parietal sulcus (IPS) and the left and right putamen.

For the first analysis of the current study, in the interest of parsimony regarding the number of areas (nodes) in our models, and given that the *current data* suggested no strong reason to assume lateralized differences in the function of the currently defined underlying network, we opted to include only the pre-SMA/SMA and the remaining left hemisphere regions as our ROIs (Figure 1d) (although see Filmer et al. 2013; Dux et al. 2006; Dux et al. 2009; Erickson et al. 2007; Erickson et al. 2005, for evidence that the brain regions supporting multitasking may be relatively more left lateralized). Upon completion of this analysis, we then sought to understand which of our conclusions might be hemisphere specific. It is worth noting that we cannot conclusively infer whether any lateralized differences are due to genuine functional hemispheric differences, or extraneous factors. However, such an analysis does provide insights into which conclusions can be drawn generally, regardless of hemisphere (and consequent decisions over the model space), and those which are hemisphere specific. To this end we repeated the analysis using the right hemisphere regions as our ROIs. We discuss the results of the left hemisphere analysis first, while referencing which findings did and did not generalise to the right hemisphere. We then present the details of the findings from the right hemisphere analysis.

### Network dynamics underlying multitasking

We sought to identify how multitasking modulates connectivity between the IPS, pre-SMA/SMA and the putamen. Although our anatomically defined mask included all of SMA, the majority (78%) of participants showed peak activity in pre-SMA, defined as coordinates rostral to the vertical commissure anterior in a probabilistic atlas based on resting state data from 12 participants (Kim *et al.*, 2010). Moreover, a visual examination of the locations of the individual peaks suggest that the remaining 22 % showed peak activity changes close to this probabilistic boundary (see Extended Figure 2d). Therefore we hereon refer to the region as pre-SMA (note: the within group percentages were also comparable; practice = 83 %, control = 73 %). To achieve this, we first applied DCM to construct hypothetical networks that could underlie the observed data. These models were then grouped into families on the basis of characteristics that addressed our key questions. This allowed us to conduct random effects family-level inference (Penny *et al.*, 2010) to determine which model characteristics were most likely, given the data. Specifically we asked; 1) which region drives inputs to the proposed network, invariant to the experimental multitasking manipulation (putamen or IPS family, Figure 2a)? and 2) does multitasking modulate striatal-cortical couplings, corticocortical couplings or both (Figure 2b)? Lastly, we conducted Bayesian Model Averaging (BMA) within the winning family to make inference over which specific parameters were modulated by multitasking (i.e. is the posterior estimate for each connection reliably different from 0?).

**Figure 2:**
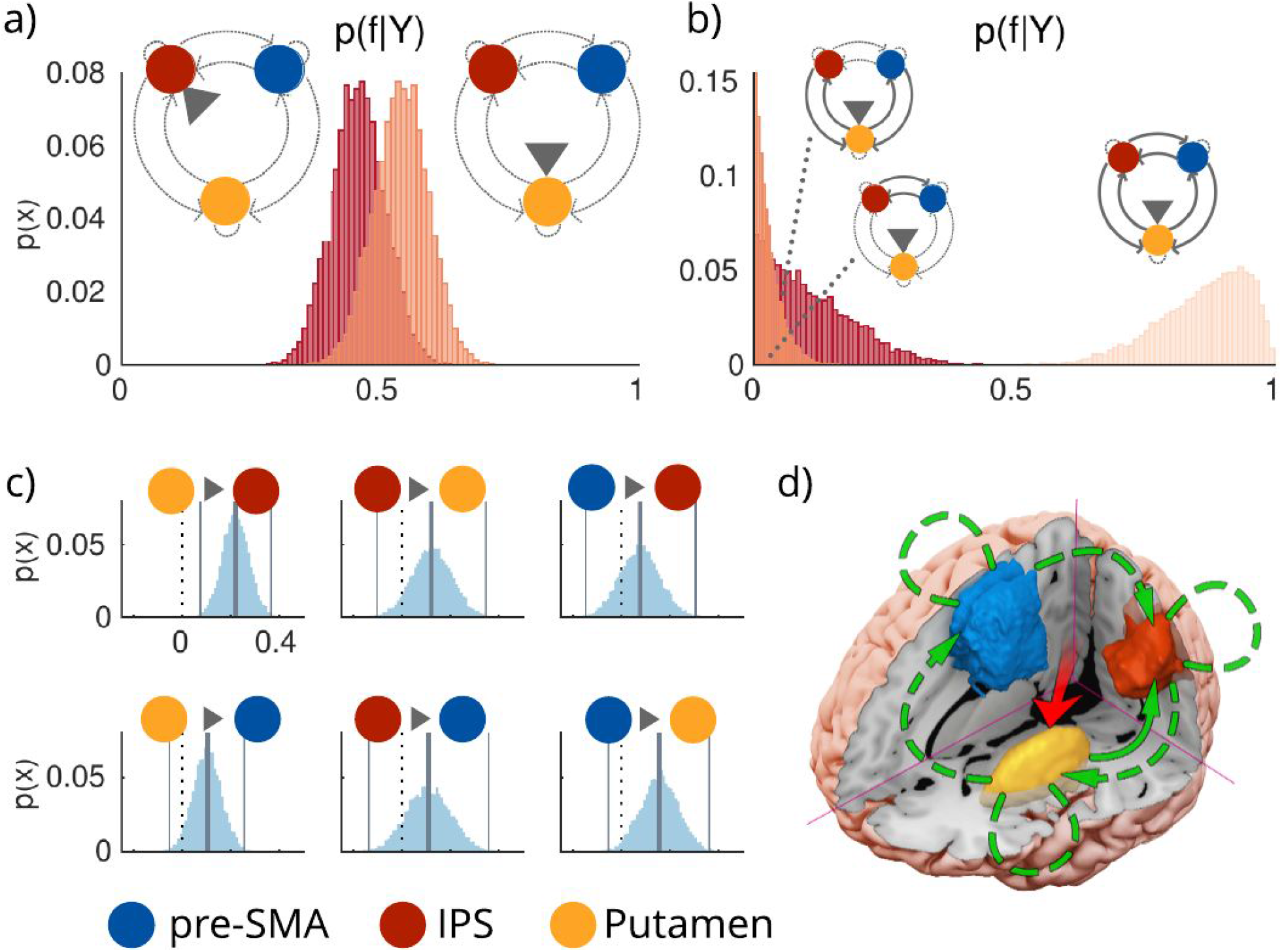
The modulatory influence of multitasking on the left hemisphere pre-SMA/IPS/Putamen network. a) Posterior probabilities over families, given the data [p(f|Y)], defined by inputs to IPS (left distribution) or Putamen (right distribution). The evidence favours driving inputs via Putamen. b) Posterior probabilities over families differing in the connections modulated by multitasking (from left to right: corticostriatal modulations, corticocortical modulations, or both). Model evidence favours both corticostriatal and corticocortical couplings. c) Posterior distributions over B parameters. Vertical lines reflect posterior means and 99th percentiles, whereas the dotted black line = 0. Multitasking reliably increased modulatory coupling from the putamen to the IPS. To view the posterior distributions that informed the removal of endogenous model connections, please check extended figure 2c. d) Proposed model for the modulatory influence of multitasking (see extended Figure 2c, for the posteriors distributions over the A parameters that motivate removal of IPS to pre-SMA and pre-SMA to Putamen). Connections drawn with a continuous line denote significantly modulated (by multitasking) connections (according to DCM.B in 2c) whereas dashed lines represent the functionally present connections (according to DCM.A in Extended Figure 2c). p(x) = probability of sample from posterior density.

The model space (Extended Figure 2a) which underpins our theoretically motivated hypothetical networks contained bidirectional endogenous connections between all three regions. Although effective connectivity can be investigated independently of anatomical connectivity, we selected this endogenous connectivity pattern given the extensive evidence for the existence of anatomical connections between the putamen, IPS and pre-SMA (Alexander, DeLong and Strick, 1986; Cavada and Goldman-Rakic, 1989; Luppino *et al.*, 1993; Wise *et al.*, 1997; Haber, 2016), as well as endogenous self-connections. As we had no a priori reason to exclude a modulatory influence of multitasking on any specific direction of coupling, we considered all 63 possible combinations of modulation (see Supplementary Figure 2b).

First we asked which region in the network received driving inputs that are modulated by multitasking demands. As the IPS shows sensitivity to sensory inputs across modalities (Grefkes and Fink, 2005; Anderson *et al.*, 2010; Vossel, Geng and Fink, 2014), and as the striatum receives sensory-inputs from both the thalamus (Alloway *et al.*, 2017) and from sensory cortices (Saint-Cyr, Ungerleider and Desimone, 1990; Reig and Silberberg, 2014; Guo *et al.*, 2018), both IPS and putamen were considered as possible candidates. Given the distribution of probability over models, it is plausible (for example) that input arrives at both the IPS and the putamen. We decided to deal with this possibility by performing Bayesian Model Averaging over the most likely models. This allows us to capture information from models that allow inputs to either the putamen or the IPS, but only under circumstances where the evidence shows that neither input should be favoured over the other. We opted to aggregate information this way as it allowed us to examine the impact of both inputs on the model evidence (separately) and retain parsimony over the model space. We therefore fit each of the 63 modulatory models twice, once allowing driving inputs to occur via the IPS, and once allowing input via the putamen (therefore, total models [M_i_] = 126). These models were grouped into two families, on the basis of their input (*IPS input* family [f_IPS_] and *putamen input* family [f_Put_]). The evidence favoured the *putamen* family (expected probability [p(f_Put_|Y)]: .54, exceedance probability (p(f_Put_|Y > f_IPS_|Y): .79, Figure 2a) relative to the *IPS* family. Therefore, the data are best explained by models where multitasking modulates driving inputs to the putamen. The winning *putamen input* family were retained for the next stage of family level comparisons. Note, for the right hemisphere, the evidence did not disambiguate between input families. Therefore we conclude that both subcortical and cortical inputs are likely to drive the network that underpins multitasking.

We then asked whether the data were better explained by models that allowed multitasking to modulate *striatal-cortical* connections, *corticocortical* connections or *all* (Figure 2b). We therefore grouped the models from the putamen input family into three groups. The *striatal-cortical family* [f_SC_] contained models that allowed multitasking to modulate any combination of the striatal-cortical connections, and none of the corticocortical connections. The *corticocortical family* [f_CC_] contained models with the opposite pattern; multitasking could modulate any pattern of corticocortical couplings and none of the striatal-cortical couplings). Finally, in the *all family*, we considered models that included modulations to both striatal-cortical and corticocortical couplings [f_ALL_]). In support of the idea that multitasking modulates striatal-cortical connectivity as well as corticocortical connections, the evidence favoured the *all* family (p(f_ALL_|Y): .86, p(f_ALL_|Y > f_SC, CC_|Y) = 1) over the *striatal-cortical* family (p(f_SC_|Y): .11) and the *corticocortical* family (p(f_CC_|Y): .03). This result was also observed for the right hemisphere.

Having determined that multitasking is indeed supported by both striatal-cortical and corticocortical couplings, we next sought to infer which specific parameters were modulated by multitasking; i.e. do we have evidence for bidirectional endogenous couplings between all regions? Or a subset of endogenous couplings? With regard to multitasking related modulations; are all couplings modulated, or a subset of striatal-cortical and corticocortical connections? To answer this we conducted BMA over the *all* family to obtain the posteriors over each of the endogenous (A) and modulatory coupling (B) parameters. We looked for A parameters to retain by testing for which posteriors showed a probability of difference (Pp) from zero that was greater than .992 (applying the Sidak adjustment for multiple comparisons). As seen in the extended Figure 2c, we retain endogenous couplings from IPS to Put, Put to IPS, Put to pre-SMA, and pre-SMA to IPS (all Pps = 1) and reject endogenous couplings from IPS to pre-SMA (Pp = .98) and pre-SMA to Put (Pp = .66). In contrast, for the right hemisphere, we retained all the endogenous connections.

We applied the same test to the B parameters and found evidence for a modulatory influence of multitasking on Put to IPS coupling (Pp = 1). Although this specific result was not found for the right hemisphere, we do find that IPS consistently shows multitasking induced coupling changes with other nodes (see the section of the right hemisphere analysis for details). Therefore we conclude that the IPS appears to be a key node in modulating information flow through the network underpinning multitasking limitations, regardless of lateralisation.

As opposed to the right hemisphere, the left multitasking-induced modulation of putamen to pre-SMA coupling did not pass our rather strict threshold of p = .98. However, it came very close. Specifically, the posterior distribution for this parameter did show reasonably strong evidence of multitask-induced modulations (Pp = .96), the variance of this distribution was more similar to the retained than the rejected coupling parameters (σ = .06 vs σ = .08), and unlike the rejected parameters, this connection showed strong evidence for the endogenous coupling (Pp = 1, Figure 2C)]. Furthermore, looking ahead to the right hemisphere analysis, we find further evidence that this coupling is modulated by multitasking demands. We therefore conclude that there is reasonable evidence that putamen to pre-SMA coupling is modulated by multitasking. We reject a modulatory influence of multitasking on the remaining parameters (all Pps <= .88).

To sum up (Figure 2d), the influence of multitasking is best explained in the left hemisphere by a network where information is propagated, via Put, to the IPS and the pre-SMA. Information is shared back to the Put via IPS, and from pre-SMA to IPS. Overall, we conclude that multitasking demands specifically increases the rate of information transfer sent from Put to pre-SMA, and between the IPS and other subcortical and cortical nodes. Hence, we can reject the idea that multitasking costs are solely due to limitations in a cortical network, rather they also reflect the taxation of information sharing between the Put and the other relevant cortical areas, namely the IPS and the pre-SMA.

### The influence of practice on the network underpinning multitasking

Next we sought to understand how practice influences the network that underpins multitasking on both single- and multi-task trials, for both the practice and control groups. For example, it may be that practice influences all the endogenous couplings in the network, or a subset of them. Furthermore, if practice only modulated a subset of couplings, would it only be striatal-cortical couplings, or corticocortical, or both? By comparing the practice group to the control group, we sought to identify which modulations are due to engagement with a multitasking regimen, and which are due to repeating the task only at the post-session (and potentially due to engagement with a practice regimen that did not include multitasking). To address these questions, we constructed DCMs that allowed practice (i.e. a pre/post session factor) to modulate all the possible combinations of couplings in the multitasking network defined above (4 possible connections, therefore *M*_*i*_ = 15, see extended Figure 3a). We then fit these DCMs separately to the single-task data and to the multitask data, concatenated across pre- to post- sessions. Comparable to above, we decided to leverage information across models (proportional to the probability of the model, see extended Figure 3b) and conducted random-effects BMA across the model space to estimate posteriors over the parameters. This method can be more robust when the model space includes larger numbers of models that share characteristics, as it helps overcome dependence on the comparison set by identifying the likely features that are common across models (Penny *et al.*, 2010). We compare the resulting posteriors over parameters to determine for each group, those which deviate reliably from zero for single-task trials, for multitask trials, and also whether they differ between groups (applying the Sidak correction for each set of comparisons).

The results from the analysis of posteriors over parameters can be seen in Figure 3. Findings that showed some generalisation in the subsequent right hemisphere analysis are as follows; for single-task trials, in the practice group, the practice factor modulated coupling from IPS to Put (Pp = .99, > .987, Sidak adjustment for multiple comparisons), which was also larger than that observed for the control group (Pp practice > control = .99). This was partially observed in the right hemisphere (see below for details). For the practice group, no other modulatory couplings achieved the criteria for significance (all Pps <= .96). For the control group, the practice factor modulated Put to pre-SMA couplings (Pp = 1, replicated in the right hemisphere, but for multitask trials), and the influence of practice was larger on this coupling for the control group than for the practice group (Pps control > practice = .99). Practice also modulated Put to IPS coupling (Pp = .99), and this modulation was larger for the control than the practice group (Pp = 1). For multitask trials, both groups showed practice related increases to modulations of the putamen to pre-SMA coupling (practice group Pp = .99, control Pp = 1, also see the right hemisphere analysis, where this was observed for the control group on multitask trials). Perhaps counterintuitively, these were larger for the control group than for the practice group (Pp control > practice = 1), note: we dissect this relationship further in the discussion. The remaining modulatory parameters and group differences did not achieve statistical significance (all Pps <= .93). Overall, practice largely influences coupling changes between putamen and IPS, and putamen and pre-SMA, and this is observed for both hemispheres (some additional coupling changes observed specifically for the right hemisphere, which are discussed below).

**Figure 3:**
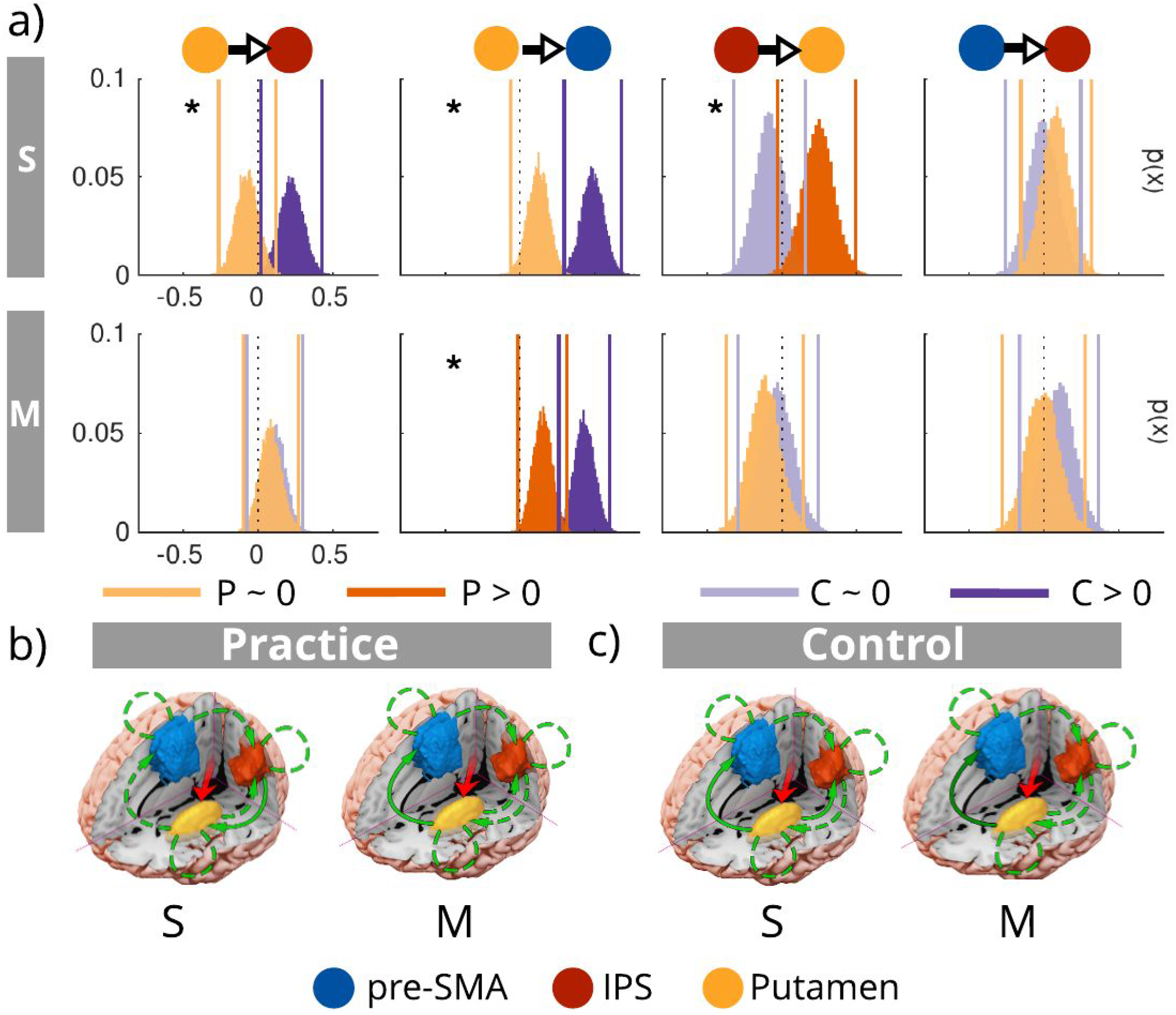
The modulatory influence of practice on the multitasking network. a) Posteriors over parameters were estimated for the practice (P, in orange) and control (C, in violet) groups for single-task trials (S) and for multitask trials (M). Posteriors that deviated reliably from 0 (>0) are in darker shades, whereas those that did not significantly deviate from 0 are in lighter shades. Stars indicate where there were statistically significant group differences. b) Proposed influences of practice on modulatory coupling within the multitasking network for single-task (S) and multitask (M) trials, for the practice and control groups. For multitask trials, the arrows are shaded to indicate the strength of the effect (i.e. the darker the arrow, the larger the modulation to that parameter between groups).

### Interrogating the right hemisphere to determine lateralization of effects

Focusing on the left hemisphere afforded a reduced model space whilst also eschewing overparameterized models. However, given that there are known hemispheric asymmetries in the brain, it is reasonable to wonder which of these results would hold, had we considered the right hemisphere. To establish which of our conclusions would have still been reached had we not used left hemisphere ROIs, we repeated the above analyses, this time using the right hemisphere (RH) data (right IPS, right Put, and pre-SMA). First we report the details of the commonalities between the left and right hemisphere analyses, and then report the specifics of the differences. Comparable to the LH analysis, practice increased the strength of putamen to pre-SMA coupling on multitask trials, however this time we observed it only for the control group (Pp = 1), and not for the practice group (Pp = .79, α_SID_= .991), although we also did not detect a difference between the two groups (Pp = .97, α_SID_= .991). Therefore, we find that multitasking influences information transfer from putamen to cortex, and that this occurs for both hemispheres.

Some lateralized differences must also be noted. For the multitasking network, in contrast to the LH model, evidence favoured neither input family (expected probability [p(f_Put_|Y)]: .5, exceedance probability (p(f_Put_|Y > f_IPS_|Y): .48), therefore the models from both input families were included in the subsequent connection family comparison. Comparable to the LH model, the evidence favoured the *all* family (p(f_ALL_|Y): .89, p(f_ALL_|Y > f_SC, CC_|Y) = 1) over the *striatal-cortical* family (p(f_SC_|Y): .09) and the *corticocortical* family (p(f_CC_|Y): .02). Unlike the LH model, posterior estimates obtained over parameters using BMA provided evidence to retain all the endogenous connections (all Pps = 1). Whereas multitasking did not modulate corticocortical connections in the LH, there was evidence that multitasking modulates RH pre-SMA -> IPS, and IPS -> pre-SMA coupling (both Pps = 1, all remaining Pps < .98). This suggests that multitasking exerts greater influence on cortical couplings in the RH than the LH, and that our conclusions are in part sensitive to the hemisphere from which we select our ROIs (see Extended Figure 5b). Importantly, and as mentioned above, the observation that multitasking modulates putamen to pre-SMA coupling is consistent across hemispheres, thus demonstrating convergent evidence that multitasking limitations stem, at least in part from modulations in information transfer between these two nodes of the network.

As the winning RH network underpinning multitasking contained all 6 endogenous connections, we considered all 63 possible combinations of modulation for the practice factor (see extended Figure 2b). Additionally, and due to the absence of evidence favouring either input, we included models that allowed inputs to the IPS and those that allowed inputs via Put (T_m_ = 126). As this constitutes the full model space, our goal was to first determine whether we could exclude families of models, prior to performing BMA in order to obtain subject level posterior estimates over parameters. We therefore conducted the same family comparisons as reported above for the multitasking network analysis. For single task trials, and in contrast to the LH, models allowing inputs to the IPS were favoured over those with inputs via Put (expected probability [p(f_IPS_|Y)]: .57, exceedance probability (p(f_IPS_|Y > f_Put_|Y): .89), whereas for multitask trials the evidence was far less conclusive (expected probability [p(f_IPS_|Y)]: .51, exceedance probability (p(f_IPS_|Y > f_Put_|Y): .58). Therefore for single trials, we retained only the f_IPS_ for the next stage of the analysis, whereas all models were retained for multitask trials. As we sought to conduct BMA over the winning family for each group separately in the next stage of the analysis, we split the practice and control group data before making model family comparisons based on connectivity patterns. For both groups, and for both single- and multi-task trials, evidence favoured the *all* family (all p(f_ALL_|Y > f_SC, CC_|Y) > .99, see Extended Figure 5c).

Examination of the posterior estimates over parameters revealed some differences in the specific couplings modulated by practice for each group. In contrast to the LH analysis, we did not find that practice modulates putamen and IPS coupling, although we did find that practice modulated IPS -> pre-SMA coupling for both the control group on single-trials (Pp = 1) and the practice group on multitask trials (Pp = 1) (see Extended figure 5d), suggesting that regardless of laterality, the IPS is a key site for practice-induced network changes. Importantly, for both the LH and RH analysis, we detect a practice related modulation on Put to pre-SMA coupling, showing that this conclusion is robust to both hemispheric and model specifications.

### Signal comparisons between left and right hemispheres

It may at first appear counterintuitive that we find some differing results between the left and right hemispheres. We know from our original analysis (Garner and Dux 2015) that our regions of interest from both hemispheres interact with our experimental factors (i.e. show a group x session x condition interaction). We have confidence in these results owing to our large sample size (N=100), and suitable corrections for multiple comparisons. As DCM models BOLD responses using a GLM, with the addition of a forward model that projects GLM parameters to a predicted BOLD response (Friston et al. 2003), we are well placed to use DCM to model our current dataset. *(Note not in the manuscript: as DCM is based on the GLM, it is amenable to examining average changes in BOLD between conditions).*

Albeit with a robust motivation, it is still useful to conduct some basic checks to inform whether the observed hemispheric differences are due to noise or are likely to be produced by genuine lateralized differences in function. One possible approach, suggested during the review process, would be to apply a simple correlation analysis to see if the observed statistical dependencies reflect what was observed for the DCM analysis. However, this unfortunately will not provide a genuine sanity check for the DCM analysis. Our concerns for applying such an analysis are as follows; correlation and DCM analyses do not test the same relationships in the data. Correlative measures test for statistical dependencies in the signal, i.e. to what extent is time series *x* associated with time series *y*? In contrast, DCM examines effective connections in the data, i.e. it asks at what rate does the theoretical neural source of time series *x* have to affect the theoretical source of time series *y*, in order to generate the best match to the observed data? Indeed this fundamental difference has been shown to yield dissociative effects. For example, autoregressive coefficients between two time series can be high, even in the absence of a direct effective connection (Friston et al. 2014; David et al. 2008). Thus, although it may be satisfying to see a correlation analysis produce the same results as the DCM, our concern is that even if this were the case, or if the correlation analysis were to show different relationships, we would be unable to draw from either observation any firm conclusions regarding the viability of the DCM analysis.

It is important to conduct sanity checks where possible, particularly when they have the capability to shed insight into how robust the current observations may be. We therefore considered what analyses we could conduct to instill such confidence (or skepticism) in the current finding of hemispheric differences in the results of the DCM analysis. As DCM serves to model the time series data from each region of interest, we reasoned that it is sensible to check the noisiness of the data underlying our current DCM analysis. We assume that a noisier time series would be more difficult to model, or indeed, may motivate overfitting of the models to the data (Lever et al. 2016). We therefore checked the ‘noisiness’ or variance of our time course data across participants, to determine whether differences between the two hemispheres could be driven by noise differences, rather than signal differences. As each time course is mean centred and adjusted for effects of no interest, we opted to take the standard deviation of the time course as a proxy of noisiness, rather than a signal to noise ratio. We therefore computed the standard deviation of each time course for each participant, region, hemisphere and DCM analysis (pre, single task pre-post, multitask pre-post). The results are presented in Figure 6. For each region, and time course and analysis, we compared the distributions between hemispheres (e.g. left IPS vs right IPS) using the z-score tests. All comparisons were not significant (*z* range: [-0.3, 0.12], all ps > .75). Therefore, the variance in the signal across participants was broadly comparable between left and right hemispheres, suggesting that the observed differences are less likely to be driven by random noise.

**Figure 6:**
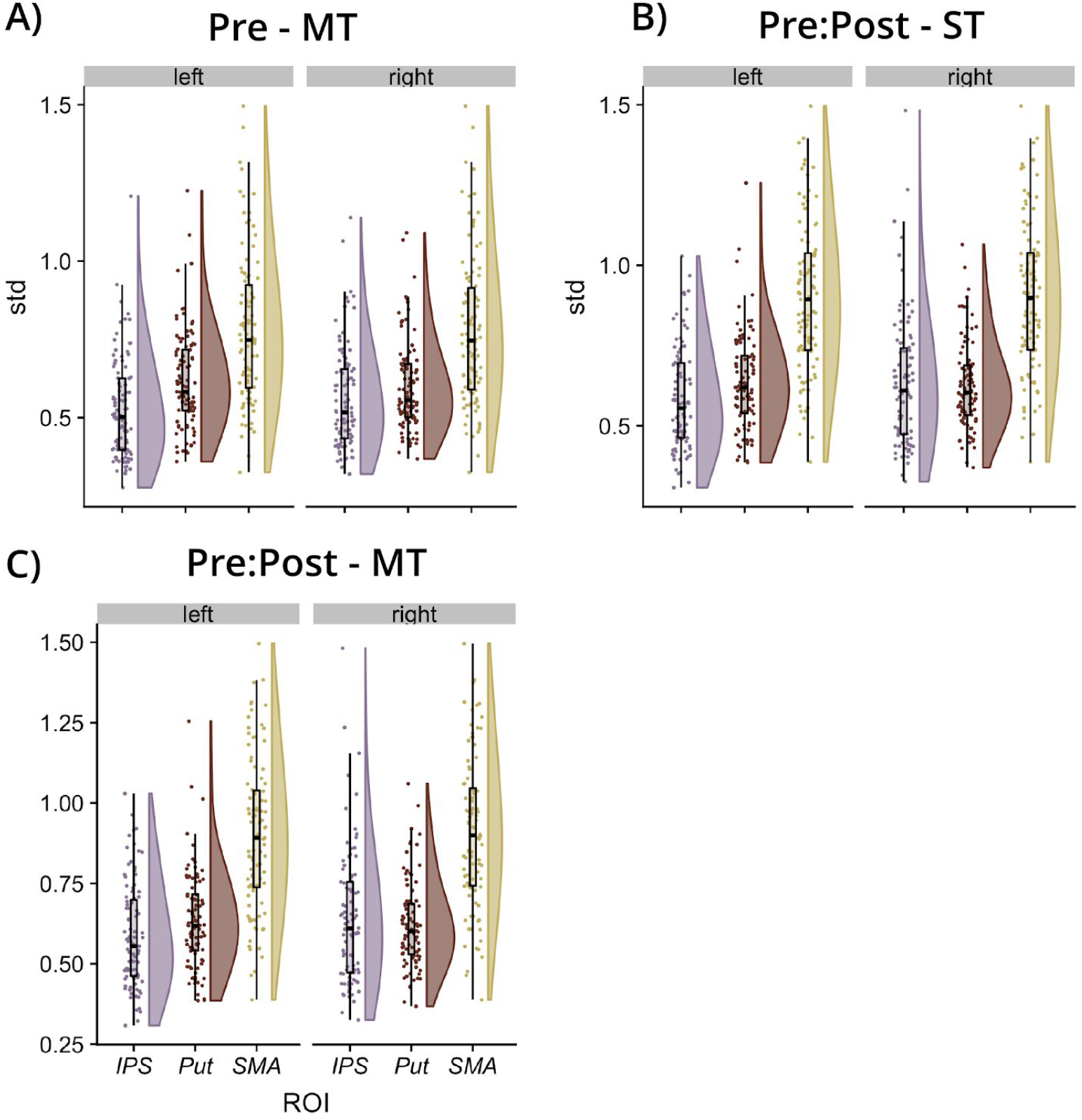
Showing the standard deviations of the mean centred time courses, across subjects, for the session 1 data (panel A), the Pre-Post single task data (panel B) and the Pre-Post multi-task data (panel B). Data are shown for each region of interest (x-axis) for the left hemisphere [left] and the right hemisphere [right]. ST = single-task, MT = multitask.

## Discussion

We sought to understand how multitasking demands modulate underlying network dynamics, and how practice changes this network to reduce multitasking costs. We adjudicated between previously hypothesised models posing that multitasking demands modulate connectivity within 1) a frontal-parietal cortical network (Jiang, 2004; Marois and Ivanoff, 2005; Dux *et al.*, 2006; Erickson *et al.*, 2007; Sigman and Dehaene, 2008; Hesselmann, Flandin and Dehaene, 2011; Tombu *et al.*, 2011; Watanabe and Funahashi, 2014; Marti, King and Dehaene, 2015), or 2) or a striatal-cortical network (Badre and Nee, 2018; Caballero, Humphries and Gurney, 2018; Yartsev *et al.*, 2018). We demonstrate evidence in keeping with the latter. Specifically, having previously identified that practice-related improvements correlate with activity changes in the pre-SMA, the IPS and the putamen (Garner and Dux, 2015), we applied DCM to ask how multitasking and practice modulates connectivity between these regions. Regardless of whether we analyse ROIs from the left or right hemisphere, we observe that multitasking consistently modulates striatal-cortical connectivity, and most consistently, information transfer from putamen to pre-SMA. Therefore, multitasking appears to modulate information sharing within a broader network than has been implied by previous studies focusing on cortical brain regions only. Rather, our results accord with models of single-task decision-making implicating a distributed striatal-cortical network. Our results build upon this work by specifically showing that attempting to multitask bilaterally increases rates of information sharing from putamen to the IPS and pre-SMA (among other modulations), and we propose that practice overcomes multitasking costs by alleviating taxation on information transfer from putamen to pre-SMA.

### Network dynamics underpinning cognitive performance in multitasking - implications

We found that during multitasking, the currently interrogated network is driven by inputs to the putamen (left and right hemisphere), and likely also the IPS (right hemisphere). While information is propagated between cortical and subcortical areas, multitasking most consistently modulates coupling from the putamen to pre-SMA. Moreover, although multitasking and practice consistently modulates the coupling between the IPS and the other nodes of the network, exactly which node appears to be dependent on the hemisphere under interrogation (putamen for left hemisphere and pre-SMA for the right hemisphere).

The IPS is assumed to contribute to the representation of stimulus-response mappings (Bunge *et al.*, 2002; Goard *et al.*, 2016; Pho *et al.*, 2018), and the pre-SMA is assumed to arbitrate between competitive representations of action-plans (Nachev *et al.*, 2007). Thus both regions potentially constitute key nodes in the cortical representation of current and upcoming stimulus-response conjunctions. Given that we observed consistent evidence that putamen to pre-SMA coupling is modulated by multitasking and practice, we propose that multitasking limitations stem, at least in part, from constraints on the rate at which the striatum can, on the basis of incoming sensory information, sufficiently excite the appropriate cortical representations of stimulus-response mappings to reach a threshold for action. This leads to the intriguing possibility that previous observations that cognitive control operations are underpinned by a frontal-parietal network (Dux *et al.*, 2009; Cole *et al.*, 2013; Duncan, 2013; Watanabe and Funahashi, 2014) may actually have been observing the cortical response to striatally mediated excitatory signals. In fact, our findings are in line with a recent application of meta-analytic connectivity modelling showing that when frontal-parietal regions associated with cognitive control operations are used as seed regions, the left and right putamen are likely to show significant co-activations across a range of sensorimotor and perceptual tasks (Camilleri *et al.*, 2018). Taken together, these data suggest that the striatum, or at least the putamen, should be included in the set of brain regions that contribute to cognitive control, at least during sensorimotor decision-making and multitasking.

It was perhaps surprising that the best network only received inputs via the putamen when the analysis was conducted on the left hemisphere data, and via both the putamen and the IPS with the right hemisphere data. However, given that a right-lateralized network incorporating the IPS has been implicated in the reorienting of attention to less probable stimuli (Vossel *et al.*, 2012), it may be that the second stimulus on multitask trials, which only occurred on one third of the trials engaged the network that responds when events demand reorientation to a new sensory input. Furthermore, although the right-hemisphere IPS showed multitasking- and practice-induced modulations of coupling with the pre-SMA, the lack of consistency between groups and hemispheres suggest that coupling activity between these nodes does not necessarily reflect the bottleneck of information processing that gives rise to multitasking costs.

### Implications of practice induced plasticity in remediating multitasking costs

Here we can both model the dynamics of the network that underpins multitasking limitations, and also identify which connections change with practice, for both single-tasks and for multitasks. By comparing this to modulations observed in the control group, we can make inroads to identifying which couplings not only correspond to multitasking limitations, but also those that may be critical in determining the extent of their presence and remediation due to practice. We interpret the control group as showing modulations that occur as a consequence of being at an earlier stage of practice (i.e. repeating the task for a second time after having practiced a regimen not expected to improve multitasking, Garner, Tombu and Dux, 2014; Garner, Lynch and Dux, 2016; Verghese *et al.*, 2017). In contrast the practice group is at a longer term stage of practice (i.e. they are repeating the task for the 5th time). It may appear counterintuitive that we observed modulations for the active control group from pre- to post- the control intervention. However, merely repeating a task without any intervening practice is sufficient to produce performance gains (Boot *et al.*, 2013). Moreover, we have observed these behavioural effects in our other practice studies (Garner, Tombu and Dux, 2014; Garner *et al.*, 2015; Verghese *et al.*, 2017) and in the current study. Importantly, here and elsewhere, practice groups show larger behavioural benefits relative to such control groups.

In light of this framework, putamen to pre-SMA coupling appears to be consistently modulated by both short- and long-term practice (the latter being more evident in the left hemisphere). We found that being at an earlier stage of practice corresponds to an increase in the rate of information transfer from putamen to pre-SMA, and that later stages are also associated with an increase of information transfer between these regions (in the left-hemisphere), but to a lesser extent than is observed for short-term practice. We interpret these results as reflecting a trajectory of practice induced changes in putamen to pre-SMA coupling. Namely, repeating a task increases putamen to pre-SMA information transfer. We speculate that extended practice results in decreased requirement for faster information transfer between these two regions, potentially reflecting decreased reliance on this pathway with extended practice. Rodent studies consistently demonstrate that when a task is novel, firing in the dorsolateral striatum corresponds to the full duration of a trial. As the behaviour becomes habitual, firing patterns transition to coincide with the beginning and end of chunked action sequences (Jog *et al.*, 1999; Barnes *et al.*, 2005; Jin and Costa, 2010; Thorn *et al.*, 2010; Smith and Graybiel, 2013). These results imply a novel physiological substrate for the amelioration of multitasking limitations; namely, the duration of information transfer from putamen to pre-SMA during task performance.

### Further considerations

It is worthwhile considering why the previous fMRI investigations into the neural sources of multitasking limitations did not implicate a role for the striatum. As far as we can observe, our sample size, and thus statistical power to observe smaller effects is substantially larger than previous efforts (Szameitat *et al.*, 2002, 2006; Jiang, 2004; Jiang, Saxe and Kanwisher, 2004; Dux *et al.*, 2006, 2009; Marois *et al.*, 2006; our N=100, previous work N range: 9-35, Stelzel *et al.*, 2006; Erickson *et al.*, 2007; Sigman and Dehaene, 2008; Borst *et al.*, 2010; Hesselmann, Flandin and Dehaene, 2011; Tombu *et al.*, 2011; Nijboer *et al.*, 2014). One fMRI multitasking study has reported increased striatal activity when there is a higher probability of short temporal overlap between tasks (Yildiz and Beste, 2015). Moreover, meta-analytic efforts into the connectivity of the frontal-parietal network during cognitive control tasks implicate the putamen (Camilleri *et al.*, 2018). Lesions of the striatum and not the cerebellum have been shown to correspond to impaired multitasking behaviours (Thoma *et al.*, 2008), and intracranial EEG has revealed that fluctuations in oscillatory ventral striatal activity predicts performance on the attentional blink task (Slagter *et al.*, 2017); a paradigm which is assumed to share overlapping limitations with those revealed by sensorimotor multitasks (Jolicoeur, 1998; Arnell and Duncan, 2002; Zylberberg *et al.*, 2010; Tombu *et al.*, 2011; Garner, Tombu and Dux, 2014; Marti, King and Dehaene, 2015). Therefore, our findings do converge with more recent efforts that do indeed implicate a role for the striatum in cognitive control. We extend these findings to demonstrate how the striatum and cortex interact to both produce and overcome multitasking limitations.

Of course, we have only examined network dynamics in a few areas of a wider system that correlates with multitasking (Garner and Dux, 2015), and we are unable to know whether we observe an interaction in these specific regions because the interaction exists nowhere else, or because the interactions are more readily observable between these regions. Indeed, we have also observed in the current dataset that the volume of the rostral dorsal lateral prefrontal cortex inversely correlates with multitasking improvements in the practice group (Verghese *et al.*, 2016) However, in this and our previous work (Garner and Dux, 2015) functional activation of the DLPFC did not meet criteria for inclusion in our analysis of the functional data. In the current study, we utilised simple sensorimotor tasks. The networks underpinning the translation of more complex sensorimotor mappings may well invoke more reliable functional activity in anterior regions of interest than we observed here (Dux *et al.*, 2006; Woolgar *et al.*, 2011; Crittenden and Duncan, 2014; Badre and Nee, 2018). Therefore, future work should determine whether more complex stimulus-response mappings would yield evidence warranting the addition of more anterior regions, such as the DLPFC, to the currently defined network.

With the current analysis, we sought to maintain parsimony by reducing the model space (and parameters) to one hemisphere, and to ascertain the findings we would have drawn regardless of which hemisphere was under interrogation. Given the bilateral nature of the observations reported in our previous work, yet the precedents in the literature for a left hemisphere bias in the networks underpinning multitasking (Filmer et al. 2013; Dux et al. 2006; Dux et al. 2009; Erickson et al. 2005; Erickson et al. 2007), we were agnostic as to whether or not we should hypothesise differences between the two hemispheres. Our analysis approach enabled us to determine that multitasking limitations and their practice-related remediation are consistently related to changes in putamen to pre-SMA coupling, regardless of which hemisphere (i.e. which subset of the reduced model space) is selected for study. It remains a little more challenging to interpret the findings that were specific for each hemisphere. Given the questions of lateralisation of function raised by the current and previous work, a principled and systematic investigation is warranted. For example, future work could simulate fMRI data in the absence of, and with genuine lateralized differences, to observe the sensitivity and robustness of DCM under these differing conditions. This simulation approach could also be applied to other degrees of freedom in the DCM analysis process. For example, we opted to use for each participant, the ROI within the anatomical region of interest, that showed strongest sensitivity to our contrast of interest. Another approach would be to apply a fixed functional ROI based on the group average. In both cases, individual outliers can be mitigated using a random effects procedure during model comparison (as we have done here). However, it remains unknown exactly how sensitive DCM analysis is to these differences in procedure. This issue is certainly not unique to our study. We are not able to test this rigorously in our own data as we do not have ground truth - i.e. we could apply a fixed ROI based on the group contrast, but as we have not generated the data that would go into each analysis, we do not exactly know how much the results of DCM analysis should differ between these analysis choices, nor exactly what these differences would mean with regards to what is the ‘correct’ model to explain the data. Once again, a principled investigation using simulated data, where the ground truth is known, is required to meaningfully address these questions but is beyond the scope of the current work.

### Conclusions

Here we asked whether multitasking limitations are better associated with activity changes in a frontal-parietal, or a striatal-cortical network. Using DCM we show evidence for the latter. Specifically, multitasking demands were associated with increased rates of coupling between the putamen and cortical sites. We interpret this as suggesting that performance decrements are due, at least in part, to a limit in the rate at which the putamen can excite appropriate cortical stimulus-response representations. Moreover, the observation that coupling strength between putman and pre-SMA is modulated with practice and that the extent of the modulation differs between the practice and control groups suggests that multitasking limits may be remediated by changes in the rate of information transfer between the putamen and the pre-SMA, that can be observed early in practice. We also suggest that modulated rates of corticostriatal information transfer gained from practice over multiple days may be a key mechanism for supporting action representations under conditions of high cognitive load. These results provide clear empirical evidence that multitasking operations may not just be mediated by a frontal-parietal network. Rather, the interface between the putamen and key cortical nodes appear to correspond to multitasking operations, and the modulation of multitasking limitations.

## Acknowledgements

This work was funded by an Australian Research Council Future Fellowship FT120100033 (PED), The University of Queensland Foundation Research Excellence Award (PED), the Australian Research Council–Special Research Initiative (ARC-SRI) Science of Learning Research Centre Grant SR120300015 (PED.), the Australian Research Council Centre of Excellence for Integrative Brain Function Grant CE140100007 (MIG), a University of Queensland Fellowship 2016000071 (MIG), and a UQ Centennial Scholarship, an International Postgraduate Research Scholarship and a Marie Sklodowska-Curie Global Fellowship (KGG). The authors thank the participants for their time, Aiman Al-Najjar, Anita Burns, Luke Hearne and Amy Taylor for assisting with data collection, and David Lloyd for assistance with results visualisation.

---

**Extended Figure 1a:**
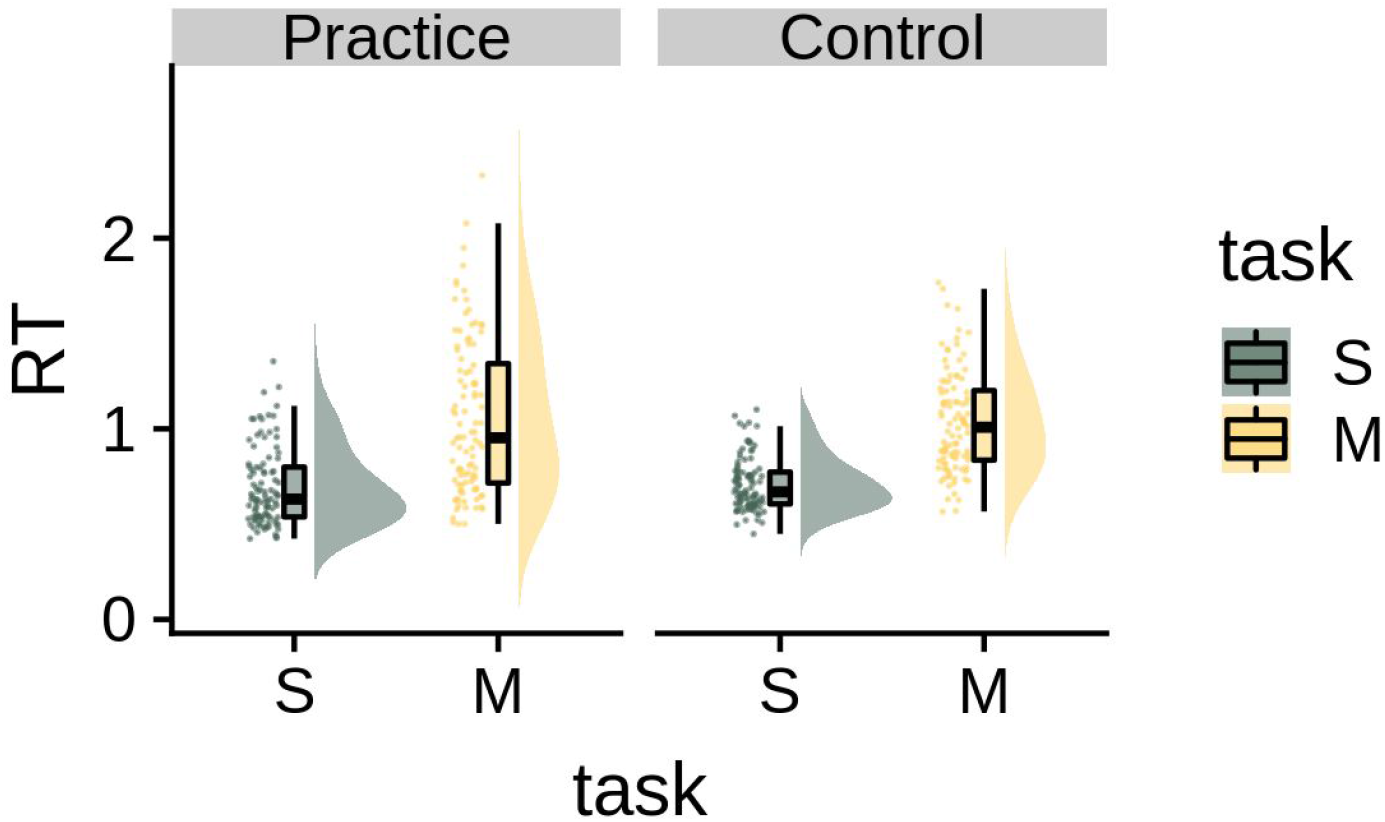
Dot, box and density plots for mean response-times (RT) for the single- (S) and multitasks (M) for the practice and the control groups at the pre-training session.

**Extended Figure 2a:**
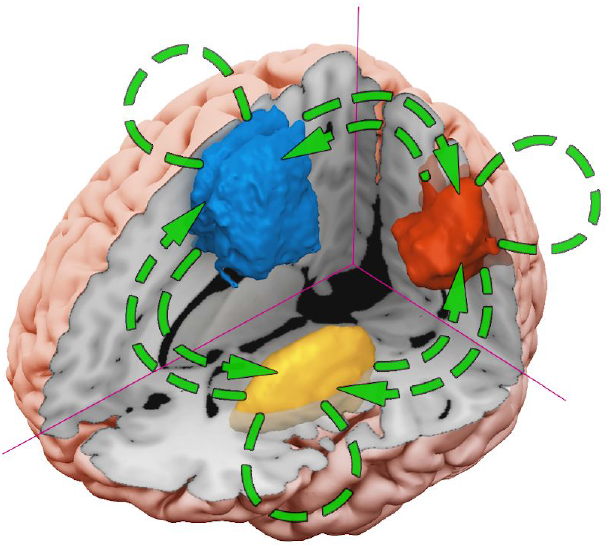
The anatomical model (DCM.A) contained bidirectional endogenous connections between all three regions, as well as endogenous self-connections.

**Extended Figure 2b:**
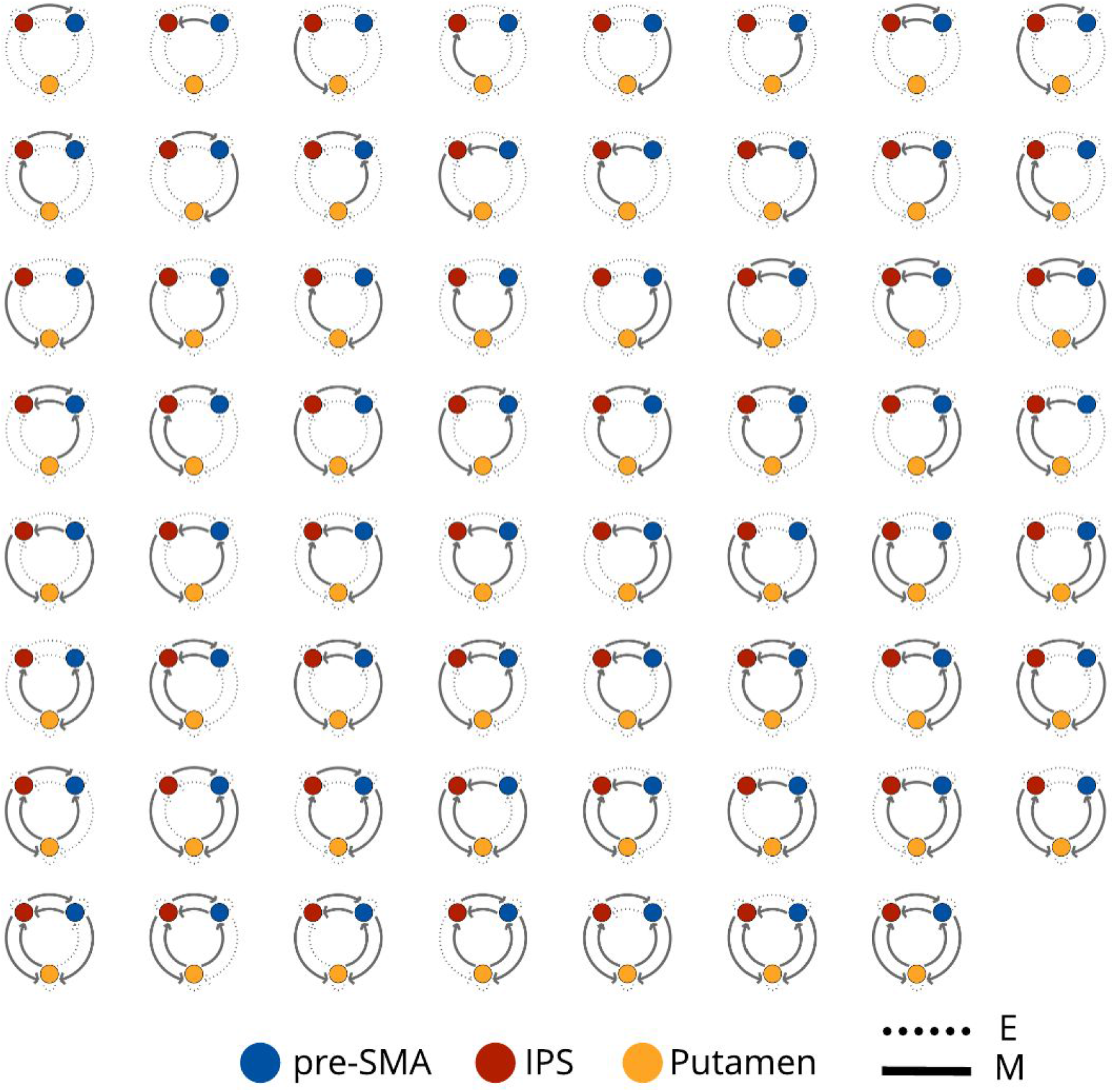
We modelled all 63 possible modulatory influences of multitasking (DCM.B). E = endogenous connections, M = modulatory connections. Each pair of regions contains bidirectional coupling.

**Extended Figure 2c:**
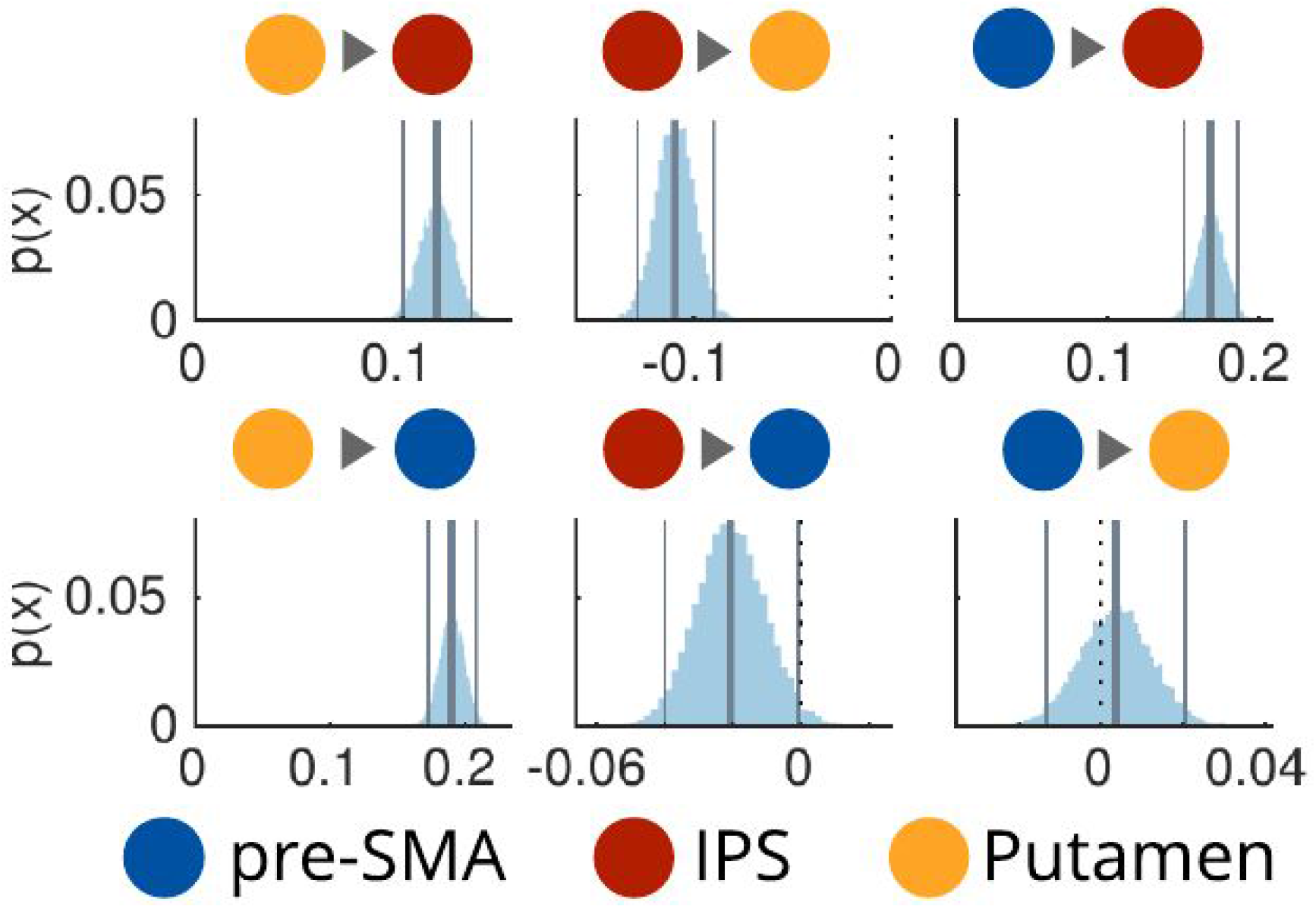
Posterior probabilities over A parameters. p(x) = probability of sample from posterior density.

**Extended Figure 2d:**
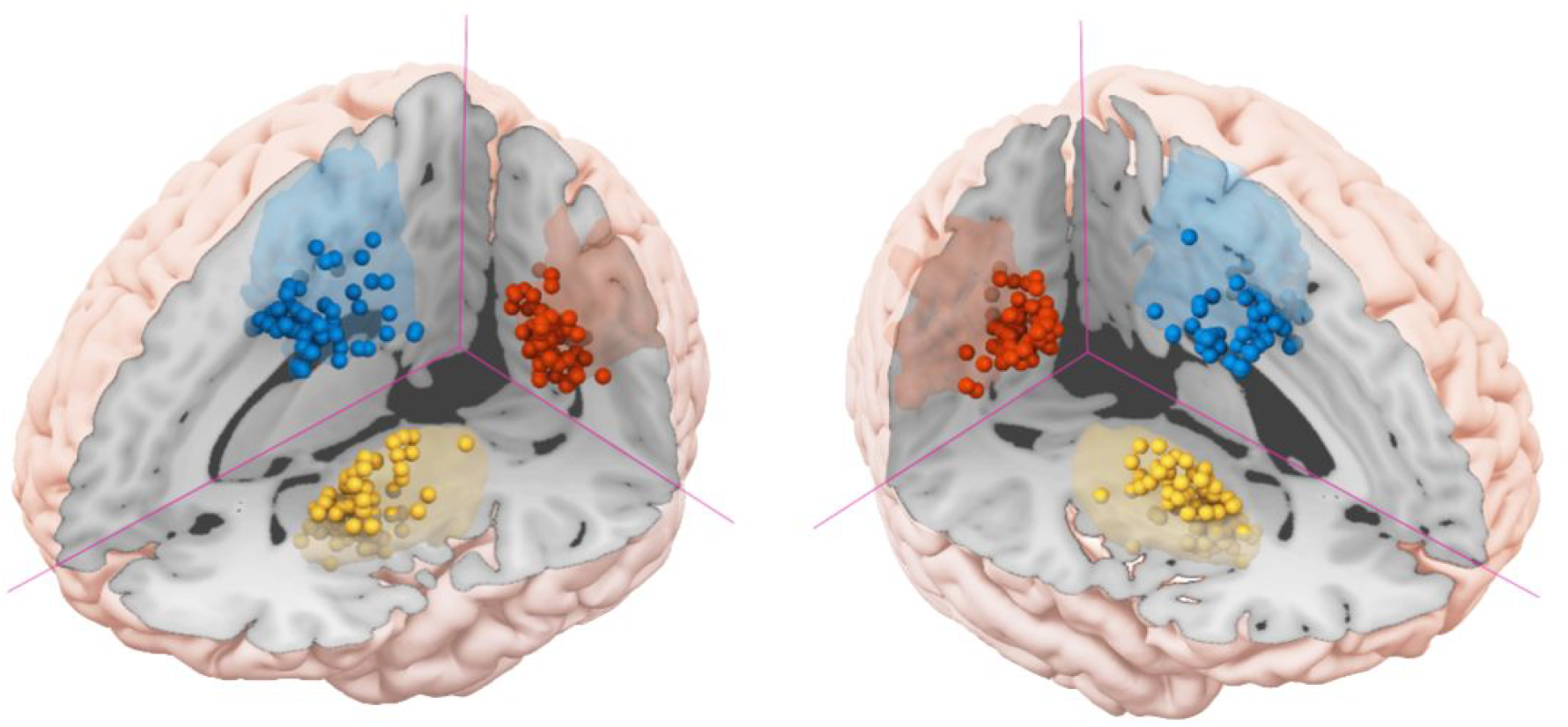
Showing the individual peaks within each region of interest for both the left and right hemisphere data (1 sphere = 1 participant).

**Supplementary Figure 3a:**
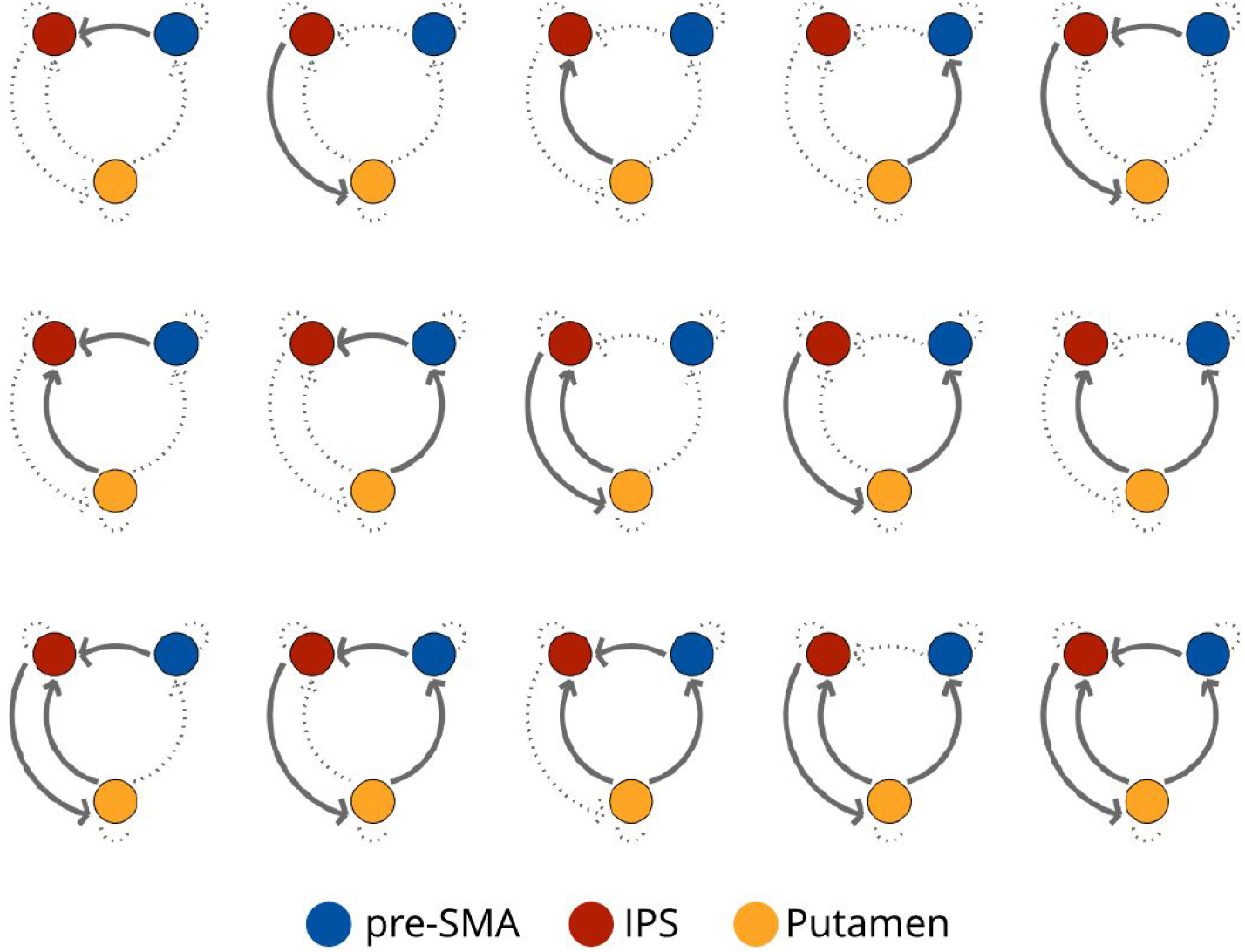
Model space considered for the modulatory influence of practice (with models *M*=1,…, 15)

**Extended Figure 3b:**
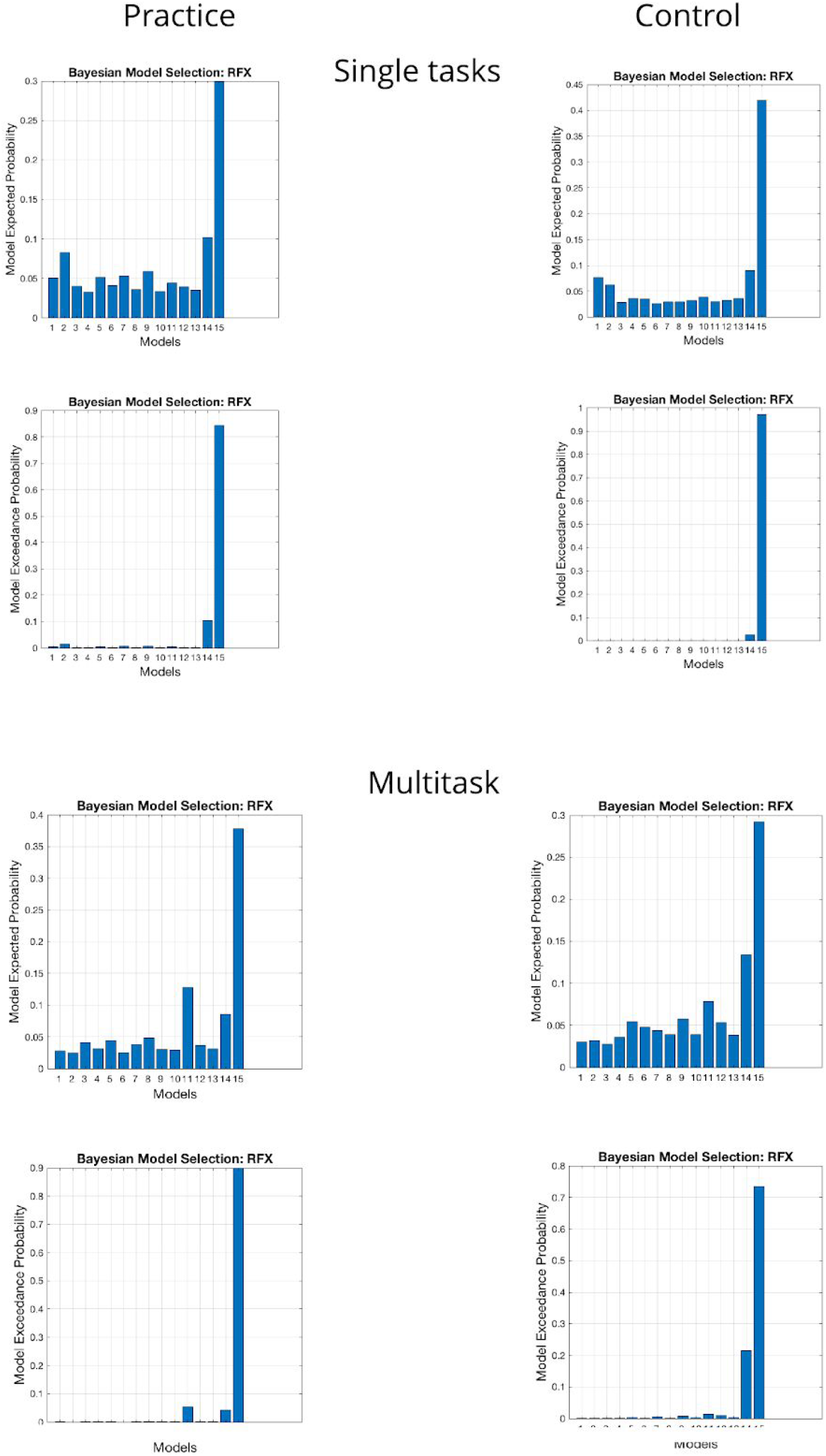
Expected and exceedance model probabilities for single-task (top 4 panels) and for multitask (bottom 4 panels) data for the practice (left column) and control (right column) groups. Models are ordered from 1-15 on the x-axis in the same order as presented in Extended Figure 3a.

**Extended Figure 5b:**
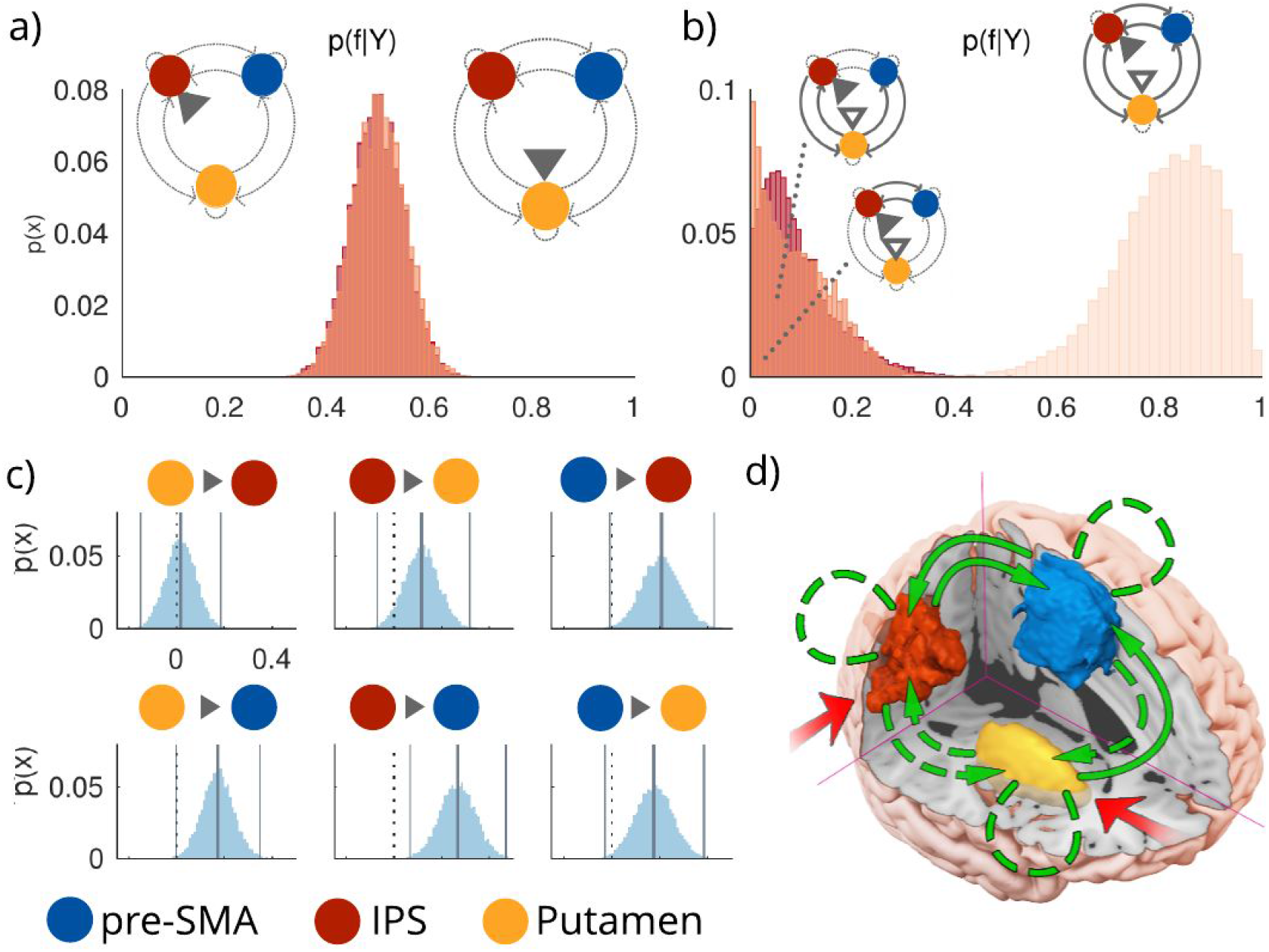
The modulatory influence of multitasking on the RH network (see control analysis section). a) Posterior probabilities over families, given the data [p(f|Y)], defined by inputs to IPS (left distribution) or Putamen (right distribution). b) Posterior probabilities over families differing in the connections modulated by multitasking (from left to right: corticostriatal modulations, corticocortical modulations, or both), averaged across families with input to either IPS or Putamen. c) Posterior distributions over B parameters. Vertical lines reflect posterior means and 99th percentiles, whereas the dotted black line = 0. d) Proposed model for the modulatory influence of multitasking in the RH. p(x) = probability of sample from posterior density.

**Supplementary Figure 5c:**
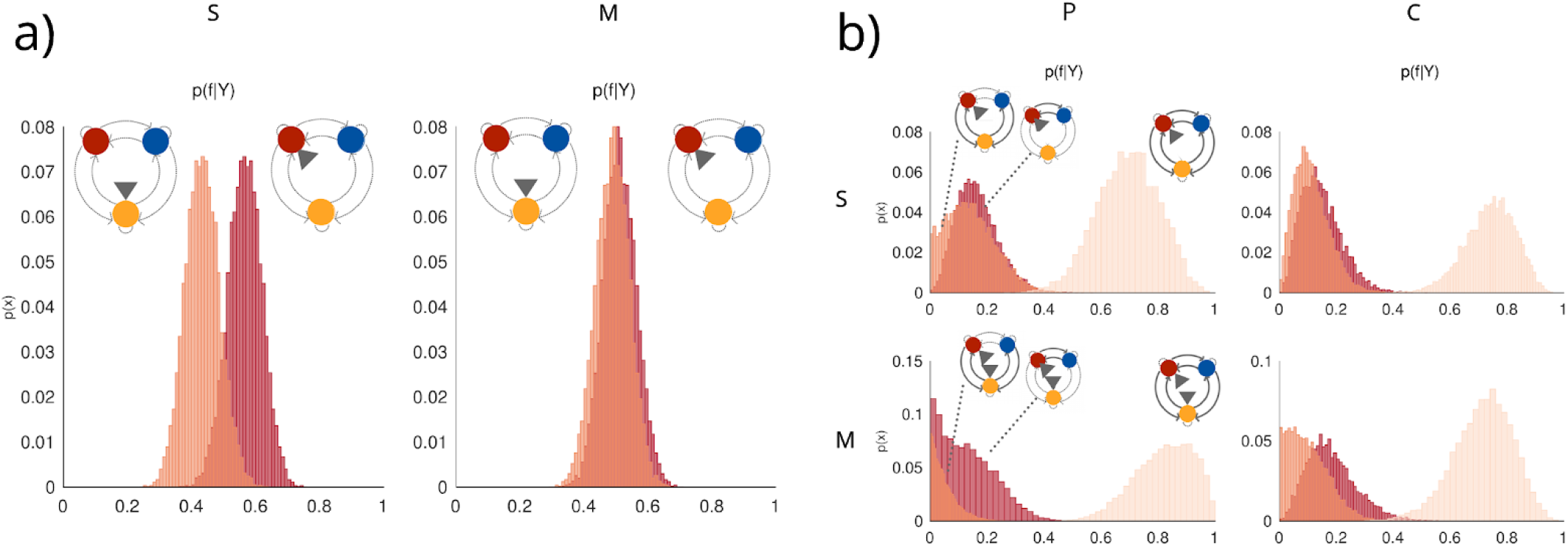
Showing model family comparisons when modelling the modulatory influence of practice. a) Posterior probabilities over families, given the data [p(f|Y)], defined by inputs to IPS (left distribution) or Putamen (right distribution) for single-task (S) or multitask (M) trials. b) Posterior probabilities over families differing in the connections modulated by multitasking (from left to right: corticostriatal modulations, corticocortical modulations, or both) for S and M trials, for the practice (P) and control (C) groups.

**Extended Figure 5d.**
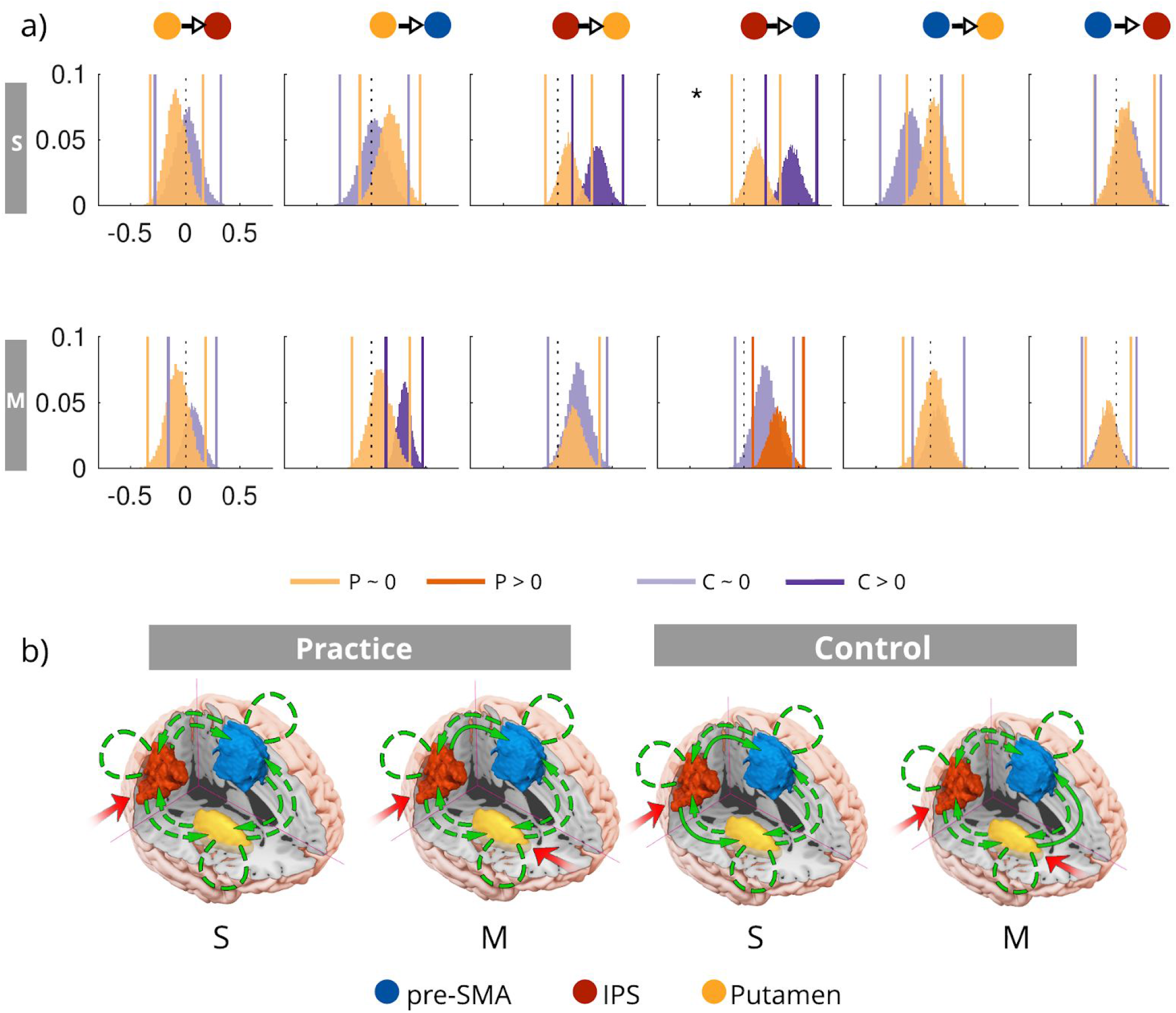
a) Showing group-level posteriors over parameters estimated for the practice (P, in orange) and control (C, in violet) groups for single-task trials (S) and for multitask trials (M). Posteriors that deviated reliably from 0 (>0) are in darker shades, whereas those that did not significantly deviate from 0 are in lighter shades. Stars indicate where there were statistically significant group differences. b) Proposed influences of practice on modulatory coupling within the multitasking network for single-task (S) and multitask (M) trials, for the practice and control groups.

## Notes

https://espace.library.uq.edu.au/view/UQ:370251

https://github.com/kel-github/multi-practice-repository

### Competing Interest Statement

The authors have declared no competing interest.

### Summary of Updates

Added a s2n analysis to check signal quality between hemispheres

